# Validating MEG source imaging of resting state oscillatory patterns with an intracranial EEG atlas

**DOI:** 10.1101/2022.10.03.510665

**Authors:** Jawata Afnan, Nicolás von Ellenrieder, Jean-Marc Lina, Giovanni Pellegrino, Giorgio Arcara, Zhengchen Cai, Tanguy Hedrich, Chifaou Abdallah, Hassan Khajehpour, Birgit Frauscher, Jean Gotman, Christophe Grova

**Affiliations:** Multimodal Functional Imaging Lab, Biomedical Engineering Department, McGill University, Montreal, Québec, Canada; Montreal Neurological Institute, Department of Neurology and Neurosurgery, McGill University, Montreal, Québec, Canada; Centre De Recherches En Mathématiques, Montreal, Québec, Canada; Electrical Engineering Department, Ecole De Technologie Supérieure, Montreal, Québec, Canada; Brain Imaging and Neural Dynamics Research Group, IRCCS San Camillo Hospital, Venice, Italy; Physics Department and PERFORM Centre, Concordia University, Montreal, Québec, Canada

**Keywords:** Intracranial EEG, Magnetoencephalography, Source imaging, Validation, Resting state, Spectral analysis

## Abstract

**Background:** Magnetoencephalography (MEG) is a widely used non-invasive tool to estimate brain activity with high temporal resolution. However, due to the ill-posed nature of the MEG source imaging (MSI) problem, the ability of MSI to identify accurately underlying brain sources along the cortical surface is still uncertain and requires validation.

**Method:** We validated the ability of MSI to estimate the background resting state activity of 45 healthy participants by comparing it to the intracranial EEG (IEEG) atlas (https://mni-open-ieegatlas.research.mcgill.ca/). First, we applied wavelet-based Maximum Entropy on the Mean (wMEM) as an MSI technique. Next, we converted MEG source maps into intracranial space, by applying a forward model to the MEG reconstructed source maps and estimated virtual IEEG (VIEEG) potentials on each IEEG channel location and quantitatively compared those with actual IEEG signals from the atlas for 38 regions of interest in the canonical frequency bands.

**Results:** The MEG spectra were more accurately estimated in the lateral regions compared to the medial regions. The regions with higher amplitude in the VIEEG than in the IEEG were more accurately recovered. In the deep regions, MEG estimated amplitudes were largely underestimated and the spectra were poorly recovered. Moreover, the MEG largely overestimated oscillatory peaks in the alpha band, especially in the anterior and deep regions. This is possibly due to higher phase synchronization of alpha oscillations over extended regions, exceeding the spatial sensitivity of IEEG but detected by MEG. Importantly, we found that MEG estimated spectra were more comparable to spectra from the IEEG atlas after the aperiodic components were removed.

**Conclusion:** This study identifies brain regions and frequencies for which MEG source analysis is likely to be reliable, a promising step towards resolving the uncertainty in recovering intracerebral activity from non-invasive MEG studies.

**Highlights:**

- Validation of MEG source imaging with intracranial EEE atlas
- Assessment of resting state human brain oscillations from healthy brain
- Adapted source imaging method, wMEM, to localize resting state oscillations
- Identified brain regions with oscillations accurately estimated by MEG
- MEG estimated spectra dominated by oscillations in the alpha band

## 1 Introduction

Neuronal oscillations are fundamental properties of brain activity and are considered to play an important role in processing and regulating neuronal communication in physiological (Giraud & Poeppel, 2012; Pellegrino et al., 2021; Voytek et al., 2010; Wang, 2010) and pathological conditions (Buzsáki et al., 2013; Hirano & Uhlhaas, 2021; Schnitzler & Gross, 2005). Electro/magneto-encephalography (EEG/MEG) are widely used non-invasive electrophysiological methods to measure neuronal activity. They provide excellent temporal resolution in the order of milliseconds, which enables to study spontaneous brain activity and oscillations in different frequency bands. Due to their non-invasive nature, EEG/MEG have been used in many studies of brain dynamics and networks, not only during well controlled tasks but also during the resting state, a state when the brain activity is spontaneous (thinking of nothing/not performing any task) (Brookes et al., 2011; Hipp et al., 2012; Keitel & Gross, 2016; Mellem et al., 2017). EEG/MEG have also been widely used as a presurgical tool for drug-resistant epilepsy and basic epilepsy research (Dalal et al., 2013; Hamandi et al., 2016; Pellegrino et al., 2018; von Ellenrieder et al., 2016). Compared to other invasive and non-invasive modalities, EEG/MEG have limited spatial resolution, since they consist in scalp recordings and source localization requires solving an ill-posed inverse problem (Darvas et al., 2004). The effects of inverse solution leakage and the challenges of localizing signals from deep brain structures are of great concern, especially when considering clinical applications such as pre-surgical planning for epilepsy (Aydin et al., 2020; Hedrich et al., 2017) and particularly while interpreting results from resting state activity due to its low signal-to-noise ratio, which is even lower for deep sources. Validation is thus necessary for non-invasive EEG/MEG techniques, to accurately interpret the results. In this study, we aimed to validate the ability of MEG source imaging to estimate resting state oscillations in healthy subjects. Due to the frequent lack of a ground truth, validation of source imaging techniques using realistic simulations are common and often useful, such as in the context of epileptic spikes (Becker et al., 2015; Chowdhury et al., 2016; Grova et al., 2006) and connectivity studies (Wang et al 2014). However, generating realistic simulations of brain activity is challenging and even more for resting state activity.

The gold standard to validate non-invasive methods is intracerebral EEG (IEEG), an invasive technique employed in some patients with epilepsy during pre-surgical evaluation. In IEEG, electrodes are placed on or into brain tissue (Jayakar et al., 2016). They are subdural grid/strip or depth electrodes (freehand or using stereoencephalography, SEEG introduced by Bancaud and Talairach in the 1950s (Enatsu & Mikuni, 2016)). IEEG thus can measure brain activity directly from the regions of interest, however, at a cost of requiring a surgical procedure to implant the electrodes and of having a limited spatial coverage. IEEG can record brain activity with excellent spatial accuracy, however validation could only be partial because of the limited spatial sampling, due to the invasiveness of the procedure.

Simultaneous recordings of EEG/MEG and IEEG provide an excellent opportunity to validate non-invasive results (De Stefano et al., 2022; Koessler et al., 2010; Pizzo et al., 2019). However, acquiring simultaneous MEG and IEEG is technically challenging (Badier et al., 2017; Dubarry et al., 2014; Kakisaka et al., 2012; Rampp et al., 2010; Santiuste et al., 2008) and not many groups have the access and technical resources to conduct such acquisitions. Also, only the patients who are candidates for epilepsy surgery undergo such invasive IEEG procedures. The implantation of intracranial electrodes is usually limited to affected regions, with a few electrodes placed in healthy regions, thus providing limited coverage of the brain. Our group developed an atlas of healthy IEEG data (Frauscher et al., 2018) at the Montreal Neurological Institute (MNI) (https://mni-open-ieegatlas.research.mcgill.ca/). This MNI IEEG atlas is generated by pooling IEEG data from 110 patients with refractory epilepsy who underwent IEEG implantation for clinical evaluation for epilepsy, only keeping the data from the electrodes implanted in healthy brain regions. With a dense coverage of all regions, this atlas provides us with the unique opportunity to study the spectral characteristics of normal brain oscillations at a group level. We took this opportunity to validate the non-invasive modality, MEG, to localize the spectral properties of the normal brain in wakefulness, in a group of healthy participants and compare those with the MNI IEEG atlas as ground truth, assuming both modalities represent the activity of the healthy brain at a group level.

We assessed MEG source imaging of resting state oscillatory patterns of healthy subjects at a group level and validated with the MNI IEEG atlas. We expect that MEG source imaging can recover the spectral patterns observed in the MNI IEEG atlas more accurately in some regions than in others. To investigate this question, we applied wavelet-based Maximum Entropy on the Mean (wMEM) (Aydin et al., 2020; Lina et al., 2012; Pellegrino et al., 2016; von Ellenrieder et al., 2016). wMEM is an EEG/MEG source imaging technique we developed and adapted to localize resting state oscillatory patterns, which proved its unique ability to recover the location and the spatial extent of the underlying oscillatory generators (Avigdor et al., 2021; Aydin et al., 2020; Pellegrino et al., 2016; von Ellenrieder et al., 2016). An original method proposed by our group (Abdallah et al., 2022; Grova et al., 2016) to estimate IEEG signals from MEG sources was then applied to support a quantitative comparison between the MNI IEEG atlas (electrical potentials) and MEG sources (cortical current densities), at the location of each IEEG electrode contact of the atlas. This is the first study to provide a group level validation with IEEG spectral characteristics across the human cortex obtained from non-invasive resting state MEG recordings in the healthy brain.

## 2 Material and methods

### 2.1 Experimental design

Our analysis pipeline is summarized in Fig 1. We used the MNI IEEG atlas as ground truth to validate MEG source imaging of resting state oscillatory patterns for healthy subjects. The MEG data were collected from 45 healthy subjects. To solve the inverse MEG problem, we applied the wavelet Maximum Entropy on the Mean (wMEM), developed by our group (Lina et al., 2012). The reconstructed MEG data along subject specific cortical surface were projected to the positions of IEEG electrodes used in the atlas to generate virtual IEEG (VIEEG) data using a method proposed by (Grova et al., 2016). To do this, the positions of the intracranial electrodes were projected from the template ICBM152 anatomy to the anatomy of each healthy subject. By applying an IEEG forward model from the sources localized along the cortical surface to all IEEG channel positions, this method allowed a quantitative comparison between the spectral properties of MEG estimated VIEEG with actual IEEG atlas for each regions of interest (ROIs) for each frequency band of interest.

**Figure 1:**
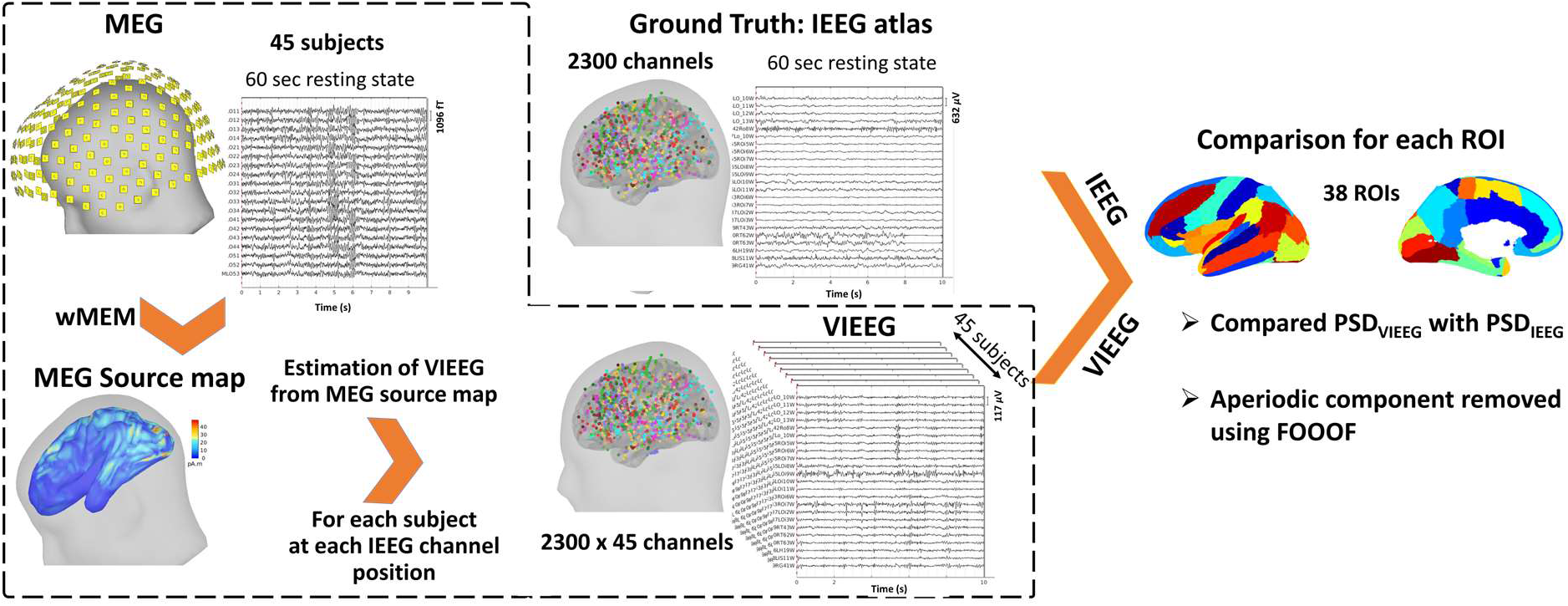
Analysis pipeline to compare the spectral properties estimated by MEG with the MNI IEEG atlas, as ground truth. **Ground truth MNI IEEG atlas** (Frauscher et al., 2018)consists of 2300 channels collected from 110 subjects with epilepsy, retaining only the healthy brain regions. For each IEEG channel, 60 seconds of resting state data during wakefulness were used. **MEG pipeline:** MEG data were collected from 45 healthy participants, each having 60 seconds of resting state data during wakefulness (only 10 seconds of data are shown in this figure). We applied wavelet-MEM (wMEM) to solve the MEG inverse problem. For each source map, we estimated virtual IEEG (VIEEG) data at each position of 2300 channels (positions obtained from the MNI IEEG atlas). We compared the spectral characteristics (spectra and oscillatory peaks) between IEEG and VIEEG for 38 ROIs (MICCAI atlas). To consider only the oscillatory components, the aperiodic components were removed from the spectra using the FOOOF toolbox (Donoghue et al., 2020).

### 2.2 Ground truth: MNI IEEG atlas

The data in the MNI IEEG atlas (Frauscher et al., 2018) were collected from 110 patients with refractory epilepsy who underwent IEEG implantation for clinical evaluation for epilepsy surgery. The key features of the data are: i) some electrodes were implanted in brain regions that turned out to be healthy and only those were retained to construct the atlas, ii) recordings were controlled with subjects having their eyes closed, and iii) electrodes were projected on the standard ICBM152 template. In the intracranial atlas, a total of 2300 channels from 110 patients (age: 31±10 Y, range: 13-62 Y, M:54) were selected. IEEG data in each patient were re-referenced to a common average reference, calculated by taking the average of 5% of the total channels exhibiting the lowest power and subtracting that value from each channel. Sixty seconds of resting state data during wakefulness were available for each of the 2300 channels. The IEEG channels in the atlas were classified into 38 regions of interest (ROIs) based on the MICCAI (Landman & Warfield, 2019) atlas. The channels from the left and the right hemispheres were considered together. The number of channels in each ROI was variable but ensured sufficient coverage of all regions: mean ± standard deviation: 60±47 channels in each ROI. More details on the data, centers, patient information, and inclusion criteria can be found in Frauscher et al. (2018).

### 2.3 Subject selection criteria for MEG

57 healthy participants who underwent MEG acquisition were included in this study (Pellegrino et al., 2022). MEG data were collected at the MEGLab of the IRCCS San Camillo Hospital in Venice, Italy. Eight minutes of resting state were acquired with eyes closed. The participants did not have any history of neurological or psychiatric disorders or any irregularity in the cycle of sleep-wakefulness. After preprocessing and sleep scoring of data, we finally included 45 participants (age: 28.67 ± 4.13 Y, range: 20-38 Y, M: 10). Of the participants, one was excluded for sleeping during the acquisition and 11 for coregistration issues such as issues with segmentation, or very noisy data.

### 2.4 MEG data acquisition

MEG data were acquired using a CTF-MEG system (VSM MedTech Systems Inc., Coquitlam, BC, Canada) with 275 axial gradiometers with a sampling rate of 1200 Hz. Bipolar electrodes were added to record Electrocardiogram (ECG) and electrooculogram (EOG). The coils were positioned on three anatomical landmarks (left and right preauricular points and nasion). These positions, along with the shape of the head of each participant were recorded with a 3D Polhemus localizer (Pellegrino et al., 2022), which were used for coregistration of MEG sensors with individual anatomical MRI of the participants.

### 2.5 Anatomical MRI, MEG-MRI co-registration and forward model estimation

For each participant, a T1-weighted-3D-TFE anatomical MRI was performed with a 3T Ingenia CX Philips scanner (Philips Medical Systems, Best, The Netherlands). The following parameters were used for MRI acquisition: [TR]=8.3 ms, [TE]=4.1 ms, flip angle=8°, acquired matrix resolution=288 × 288, slice thickness=0.87 mm) (Pellegrino, 2022). Freesurfer (Dale et al., 1999) was used for subsequent brain segmentation and reconstruction of the white/gray matter interface. The coregistration of MEG sensors with anatomical MRI was performed in Brainstorm (Tadel et al., 2011), applying a surface fitting between the head shape from MRI and the positions of coils and head shape recorded using 3D Polhemus during MEG acquisition. We considered the cortical mesh of the mid layer which is equidistant from the white and grey matter interface as source space, consisting of around 8000 vertices. The forward model was computed using OpenMEEG software (Gramfort et al., 2010; Kybic et al., 2005) implemented in Brainstorm (Tadel et al., 2011). We used a 3-layer Boundary Element model (BEM) consisting of brain, skull, and scalp surfaces with conductivity values of 0.33, 0.0165, and 0.33 S m^-1^, respectively (Zhang et al., 2006).

### 2.6 MEG data preprocessing

MEG preprocessing was performed with Brainstorm software (Tadel et al., 2011). Preprocessing of MEG data included (i) filtering within the 0.5-80 Hz band, (ii) applying a notch filter at 50 Hz, (iii) downsampling to 200 Hz, (iv) applying third-order spatial gradient noise correction and (v) removal of cardiac and eye movement artifacts using Signal Space Projection (SSP) (Uusitalo & Ilmoniemi, 1997) routine available in Brainstorm. A sixty-second segment was extracted for each subject, continuous or concatenated (minimum length of the continuous segment: 10 seconds), where no artifact was visibly present, ensuring with an EEG expert that the subject was awake during this segment. To assess the data for sleep score, some scalp EEG channels were provided (F4, C4, O2, Ref left mastoid, Ground left shoulder).

### 2.7 MEG Source imaging using wavelet Maximum Entropy on the Mean (wMEM)

The MEG inverse problem was solved using the Maximum Entropy on the Mean (MEM) (Amblard et al., 2004), which we carefully validated in the context of EEG/MEG source imaging (Chowdhury et al., 2013). The key feature of this framework is a spatial prior model, assuming that brain activity is organized within cortical parcels. MEM is a Bayesian framework, where the activity of every parcel is tuned by the probability of activation of a hidden state variable. When the parcel is active, a Gaussian prior is assumed to model a priori the activity within the parcel. Starting from such a prior “reference” distribution, inference to ensure data fit is then obtained using entropic techniques. As a result, MEM is able to either switch off or switch on the corresponding parcels during the localization process, while allowing local contrast along the cortical surface within the active parcels. Parcellation of the whole cortical surface (K~228 parcels) and initialization of the probability of being active were obtained using a data driven approach, based on a Multivariate Source Pre-localization (MSP) method (Mattout et al., 2005), a projection technique allowing to define the probability of every source to contribute to the data. The MEM specific prior model, using the entropic technique to fit the data, allows accurate localization of the underlying generators together with their spatial extent, as previously demonstrated by our studies for the standard version of MEM (cMEM) (Abdallah et al., 2022; Chowdhury et al., 2016; Chowdhury et al., 2013; Grova et al., 2016; Heers et al., 2016), as well as the wavelet-based extension of MEM (wMEM) (Lina et al., 2012). wMEM was specifically designed to localize brain oscillatory patterns. wMEM applies a discrete wavelet transformation (Daubechies wavelets) to characterize the oscillatory patterns in the data before applying the MEM solver (Lina et al., 2012). We validated wMEM for localizing oscillatory patterns at seizure onset (Pellegrino et al., 2016), interictal bursts of high frequency oscillations (Avigdor et al., 2021; von Ellenrieder et al., 2016) and MEG resting state fluctuations (Aydin et al., 2020). Both wMEM and cMEM implementations are available within the Brain Entropy in space and time (Best) plugin of Brainstorm software (https://neuroimage.usc.edu/brainstorm/Tutorials/TutBEst/). To compute the sensor level noise covariance matrix from ongoing resting state data, we used a synthetic baseline generated from the segment of the signal of interest, by resampling the phase of the Fourier transform of the signal (Prichard & Theiler, 1994). This baseline preserves the spectrum of the signal while destroying the temporal coherence. We incorporated a few changes in standard wMEM implemented in Brainstorm, to localize specifically oscillatory patterns in resting state data, as described in the Appendix.

### 2.8 Estimation of virtual IEEG (VIEEG) data from the MEG source map

MEG and IEEG are two different modalities each measuring brain activity in different units, MEG measurements after source imaging are estimated current densities (in nanoAmpere-meters), whereas IEEG measurements are electrical potentials in μVolts. To allow quantitative comparison between these two modalities, we converted MEG reconstructed source maps into IEEG channel space, by estimating corresponding IEEG potentials that would correspond to those MEG sources on each electrode contact (channel) of the atlas (Abdallah et al., 2022; Grova et al., 2016). To do so, we first localized the position of all channels of the atlas within the native MRI referential system of all healthy subjects from whom we analyzed MEG data. Co-registration between anatomical MRI of each subject and the ICBM152 template where the atlas is defined was obtained using Minctracc program (Collins et al., 1994). This is obtained in three steps: (1) estimation of a linear registration to account for the linear part of the transformation (using bestlinreg_s tool), (2) estimation of a non-linear transformation to account for the variability between the two maps (using minctracc tool); (3) application of the resulting non-linear transformation to the coordinates of the electrode contacts of MNI IEEG atlas, to convert them from the ICBM152 anatomy to the anatomy of each healthy subject.

Then, for each subject, to estimate the virtual IEEG potentials from MEG estimated current density, J_MSI_, we calculated a subject specific IEEG forward model, G_IEEG_ that estimates the influence of each dipolar source of the cortical surface on each IEEG channel (Grova et al., 2016). Since we did not intend to solve the inverse problem of source localization from IEEG data, we used a simplified IEEG forward model G_IEEG_ assuming an infinite volume conductor characterized by a conductivity *σ* of 0.25 S.m^-1^. For a total number of IEEG contacts *c*, (*c* = 2300) and *n* number of cortical sources (*n* = 8000). G_IEEG_ is a *c* x *n* matrix that estimates the electrical potential located at each IEEG electrode *i* (*i=1, 2…c*) corresponding to an equivalent current dipole of unit activity located on the vertex *S_j_* and oriented along 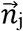, normal to the cortical surface (*j = 1,2,….n*), calculated as:

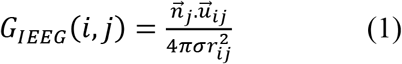

where 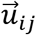 is a unit vector oriented from the source *S_j_* to the IEEG contact *i* and *r_ij_* is the Euclidean distance between *S_j_* and contact *i*. To avoid numerical instabilities, when the sources on the cortical surface were too close to the IEEG contacts (*r_ij_* < 3 mm), the distance *r_ij_* was set to 3 mm instead, keeping the orientation 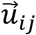. Finally, we applied the IEEG forward model, G_IEEG_ to the MEG reconstructed source map (J_MSI_) to estimate IEEG potentials on each IEEG channel, VIEEG as:

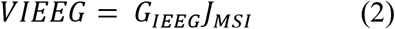

Here, we applied a simplified IEEG forward model because Cosandier-Rimélé et al. (2007) showed that it could estimate accurately real IEEG measurements. Moreover, von Ellenrieder et al. (2012) showed that the use of finite-element models considering the actual size and the shape of the IEEG electrodes had almost no influence on local electrical potentials at 2 mm from the electrodes.

As for each source map, we estimated VIEEG for each IEEG channel in the atlas, we generated more VIEEG channels compared to the MNI IEEG atlas (2300 channels in the IEEG atlas vs 2300 × 45 channels in VIEEG).

### 2.9 Frequency specific brain maps of relative power

For each of the IEEG and VIEEG channels, the power spectral density (PSD) was estimated using Welch’s method (Time duration: 0-60 seconds, 2s sliding Hamming windows, overlap: 50%). For each channel, a relative PSD was obtained by dividing each PSD value by the total power across the whole frequency range. The group average of relative PSD was calculated across all channels within a ROI and all frequency bins in each frequency band of interest: δ (0.5-4Hz), θ (4-8Hz), α (8-13Hz), β (13-30Hz), and γ (30-80Hz).

We also applied the depth weighted minimum norm estimate (MNE) as another standard source imaging method to obtain VIEEG_MNE_ to compare with IEEG and VIEEG_MEM_ (VIEEG estimated using wMEM). To calculate the noise covariance for depth weighted MNE, we used 2 seconds of resting state data from each subject. To solve MNE, we estimated the regularization hypermeter λ by using the signal-to-noise ratio (SNR) of the data, as *λ=1/SNR^2^*, with the SNR arbitrarily set to 3 (default value in Brainstorm). We used the term VIEEG_MEM_ to differentiate from VIEEG_MNE_ only for the analysis of relative PSD. For the rest of the document, we used the notation VIEEG instead of VIEEG_MEM_.

To compare the relative PSD before and after applying the conversion from MEG source maps into virtual intracranial space, we also calculated the relative PSD after MEG source imaging directly along the cortical surface (Fig S2). To obtain a group average of relative PSD for 45 subjects, we followed the approach described in Niso et al. (2019). We first projected individual relative PSD to a default template, ICBM152 (Fonov et al., 2009) and then obtained a group average of 45 relative PSD in each frequency band of interest.

### 2.10 Analysis of spectral oscillatory components

In this study, we applied FOOOF (Fitting Oscillations & One-Over-F) (Donoghue et al., 2020), a commonly used algorithm (Huang et al., 2021; Mahjoory et al., 2020; Ramsay et al., 2021; Senoussi et al., 2022; Wiesman et al., 2022) to separate periodic components from the aperiodic components of the spectra by parameterizing the power spectra as a composition of these two components. Although the aperiodic components are also of much interest and recently gained attention (Bódizs et al., 2021; He, 2014; Ostlund et al., 2021; Ouyang et al., 2020; Schaworonkow & Voytek, 2021; Wilkinson & Nelson, 2021), our aim in this study was to compare and validate the periodic components only.

For each of the IEEG and VIEEG channels, we decomposed the spectra into periodic and aperiodic components using the FOOOF toolbox (MATLAB version) (Donoghue et al., 2020). The following FOOOF parameters were used: frequency range = 0.5–80 Hz; peak type: Gaussian; peak width limits (minimum bandwidth, maximum bandwidth) = 1 – 8 Hz; maximum number of peaks = 8; peak threshold: 3.0 dB; proximity threshold = 2 SD; aperiodic mode: *knee.* As we concentrated only on the rhythmic activities of the spectra, we subtracted (in the log-log scale) the aperiodic component from the raw PSD. The remaining oscillatory component of the spectra (PSD_IEEG_ and PSD_VIEEG_) was considered for further analysis and comparison between IEEG and VIEEG. We also identified the oscillatory peaks during the process of finding aperiodic components.

### 2.11 Comparison of VIEEG spectra with IEEG

For each ROI, we calculated the median of PSD_IEEG_ across all available channels within the ROI (N_ROI_). The median of PSD_VIEEG_ for each ROI was obtained across a total number of channels = N_ROI_ x *Number of healthy subjects.* The overlap between PSD_IEEG_ and PSD_VIEEG_ was calculated for each frequency bin as:

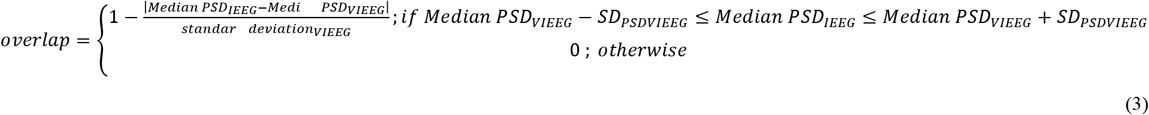

We calculated the *average overlap* for each ROI before and after removing the aperiodic components of the spectra. *Average overlap* quantifies the distance between *median PSD_VIEEG_* and *median PSD_IEEG_* at each frequency bin. The value of this metric ranges between 0 and 1. If the median of PSD_VIEEG_ perfectly coincides with the median of PSD_IEEG_ at a specific frequency, the *overlap* is 1, and if the median of PSD_IEEG_ is greater or less than one standard deviation of PSD_VIEEG_, the *overlap* is zero at that frequency. We obtained *average overlap* across all the frequency bins within each frequency band of interest: δ (0.5-4Hz), θ (4-8Hz), α (8-13Hz), β (13-30Hz), and γ (30-80Hz).

### 2.12 Comparison of VIEEG with IEEG in terms of peak frequency

We also compared the oscillatory peaks between the MEG estimated VIEEG and the MNI IEEG atlas. Using FOOOF, we identified all oscillatory peaks in each IEEG and VIEEG channel. For each ROI, the number of channels (out of the total number in each ROI N_ROI_) exhibiting an oscillatory peak in a specific frequency band of interest was calculated for the IEEG and VIEEG of each subject. Then we calculated the percentage difference of the number of channels exhibiting peak in a specific frequency band as:

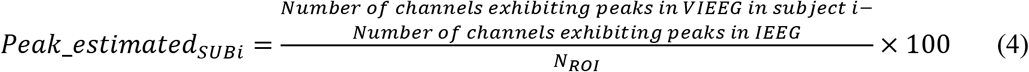

This measure was obtained for each ROI and each frequency band for each subject *i*. To obtain a group level estimation of channels exhibiting peaks per ROI per frequency band, we calculated the median of *Peak_estimated_SUBi_* over 45 subjects. This metric is a percentage assessing the overestimation or underestimation of MEG estimated VIEEG channels showing oscillatory peaks, when compared to IEEG. The value of the Median (*Peak_estimated_SUBi_*) ranges from −100% to 100%. For a particular ROI and frequency band, Median *(Peak_estimated_SUBi_)* = +100% indicates that all the VIEEG channels in that ROI (N_ROI_) showed peaks in that frequency band, whereas no peak was identified in any of the IEEG channels in that ROI and frequency band. We called it a 100% overestimation of peaks by MEG estimated VIEEG in that ROI. On the contrary, a −100% estimation (underestimation) is obtained when all the IEEG channels in a ROI exhibit peaks, but VIEEG fails to identify any peak in that ROI. The peaks are well estimated by VIEEG if the Median *(Peak_estimated_SUBi_)* is close to zero.

## 3 Results

### 3.1 Frequency specific brain maps of relative power

In Fig 2, the group average of relative PSD is plotted for 38 ROIs for IEEG, and MEG estimated VIEEG. For each frequency band in Fig 2, we used a common color bar for all modalities, thus highlighting how much MEG could estimate relative power when compared to IEEG. Fig S1 shows another representation of the same data, the color bar ranging from minimum to maximum value for each modality in each frequency band. Fig S1 highlights the regions exhibiting the strongest activation of average relative power within each modality.

**Figure 2:**
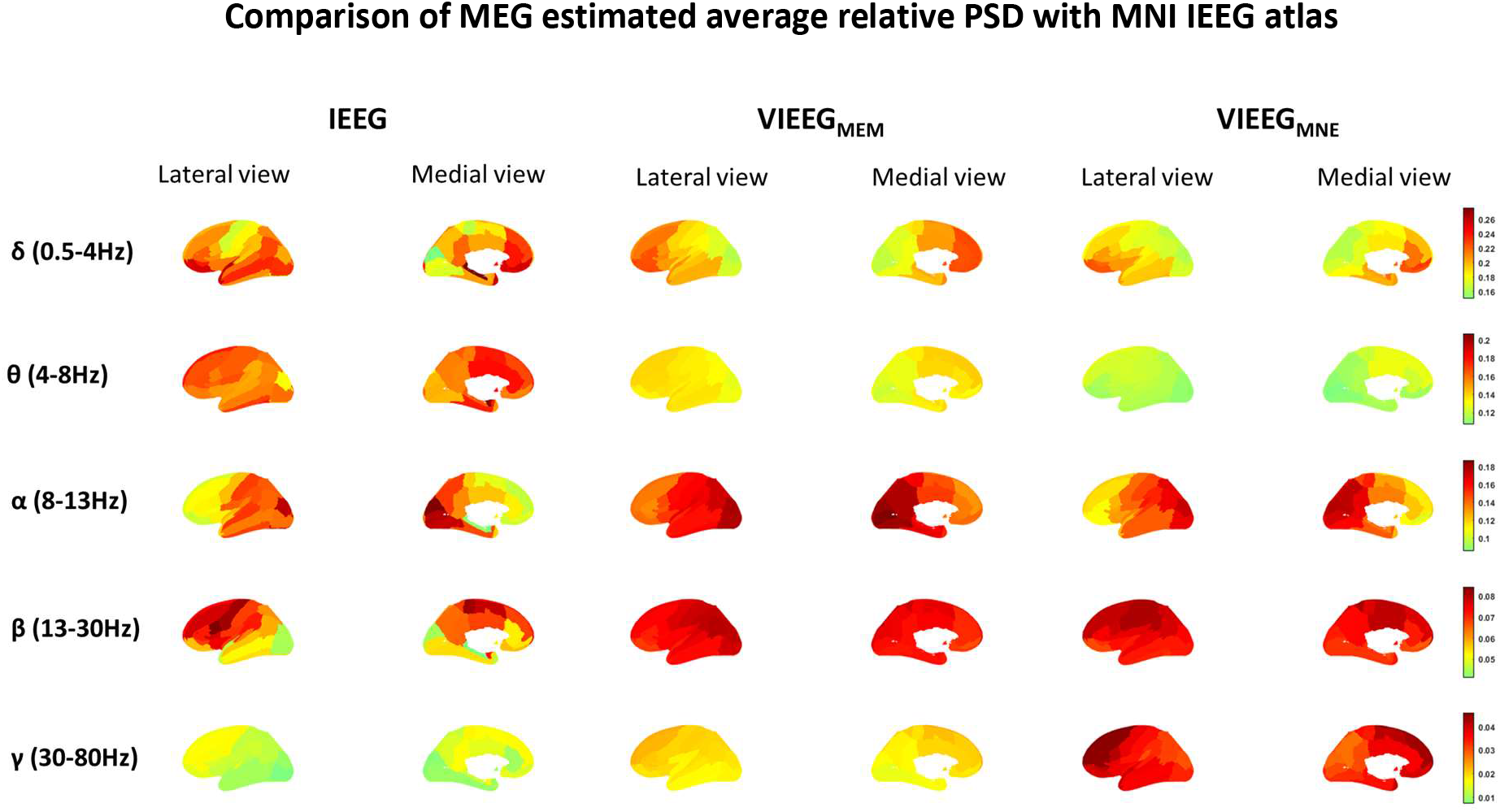
Group average of relative PSD values across each frequency band and over all the available channels in each ROI of the IEEG atlas, the ground truth, the MEG estimated VIEEG using wavelet-MEM (wMEM) method VIEEG_MEM_, and minimum norm method estimate VIEEG_MNE_. The number of channels in each ROI *(N_ROI_)* in IEEG varies. For each ROI, VIEEG was estimated for *45 x N_ROI_* channels, where the total number of subjects is 45. The relative PSD for each channel is calculated as the ratio of the power of the signal in each frequency bin relative to the total power of the signal. Relative PSD value for a channel range between 0 and 1. The color bar ranges from minimum to maximum value among IEEG, VIEEG_MEM_,and VIEEG_MNE_ in each frequency band, such that the scale is the same for all modalities in a given band.

Overall, similar patterns of power distribution were observed between IEEG and MEG estimated VIEEG (VIEEG_MEM_), such as high power in delta, theta, and gamma bands in the anterior ROIs, and high power in the alpha band in posterior ROIs. Beta power was strong around the primary and supplementary motor regions in both IEEG and VIEEG. However, for all frequency bands in Fig 2, the relative PSD in each IEEG ROI was very distinct, showing important contrasts and a larger range from strongest to weakest activity among ROIs, whereas MEG estimated VIEEG maps were smoother among the neighboring ROIs spanning a smaller range of activity. This was evident especially in deeper regions such as the hippocampus and amygdala, which were very distinguished in IEEG showing very strong or weak activity, whereas these regions showed smoother activation in VIEEG, almost undistinguishable from the neighboring ROIs. We also observed high delta power and weak beta power in IEEG in lateral posterior ROIs, which were not well estimated by VIEEG.

The MEG estimated VIEEG using depth weighted MNE, VIEEG_MNE_, also showed similar patterns of power distribution as estimated by VIEEG_MEM_. Compared to IEEG, VIEEG_MNE_ exhibited an underestimation in the theta band and an overestimation in the gamma band. In contrast, VIEEG_MEM_ power in the theta and gamma bands was in a similar range as IEEG. The relative power estimated by wMEM and depth weighted MNE on the cortical surface (Fig S2) also showed similar patterns in all frequency bands, except for gamma (more details in the supplementary material).

### 3.2 Analysis of spectral oscillatory components

In Fig 3, for four typical example ROIs selected at different depths (two in the lateral and two in the medial side), we show the decomposition of spectra into periodic and aperiodic components for IEEG and VIEEG. The spectra are plotted as median ± standard deviation across all available channels within a ROI, *N_ROI_* channels for IEEG, and the *number of healthy subjects × N_ROI_* channels for VIEEG. Fig 3 also shows the probability histogram of all identified oscillatory peaks in the ROI for VIEEG and IEEG. For comparison between the VIEEG and IEEG spectra, we only considered the periodic components of the spectra, after removing the aperiodic components from the original spectra. The comparison between IEEG and VIEEG spectra is discussed in the following sections.

**Figure 3:**
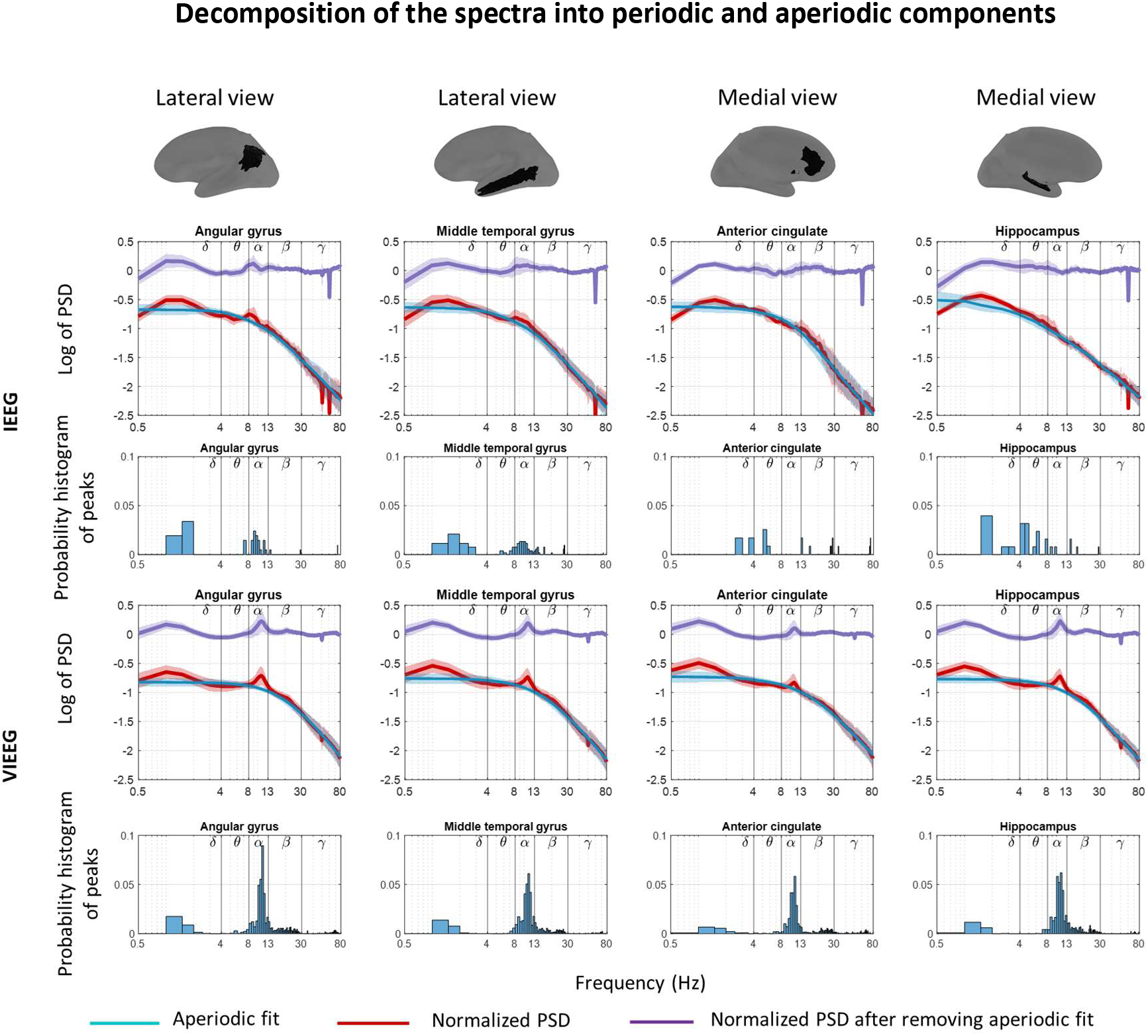
Decomposition of spectra into periodic and aperiodic components shown in a few example ROIs for IEEG and VIEEG. For both IEEG and VIEEG, the top panel shows the normalized PSD before and after removing the aperiodic component and the bottom panel shows the probability histogram of identified peaks in *δ* (0.5-4Hz), *ϑ* (4-8Hz), *α* (8-13Hz), *β* (13-30Hz) and *γ* (30-80Hz). The aperiodic fits and oscillatory peaks are identified using the FOOOF toolbox (Donoghue et al., 2020).

### 3.3 Comparison of VIEEG spectra with IEEG

We show the comparison between IEEG and VIEEG spectra for four example ROIs (two in the lateral and two in the medial side) in Fig 4, before and after removing aperiodic components. We selected these four ROIs based on how MEG estimated the spectra compared to IEEG, quantified by the metric *average overlap* (performance worsening from left to right, after removing the aperiodic components). The figure shows the value of *average overlap* quantified for each frequency band for a ROI. For all four ROIs, the *average overlap* improved after removing the aperiodic components.

**Figure 4:**
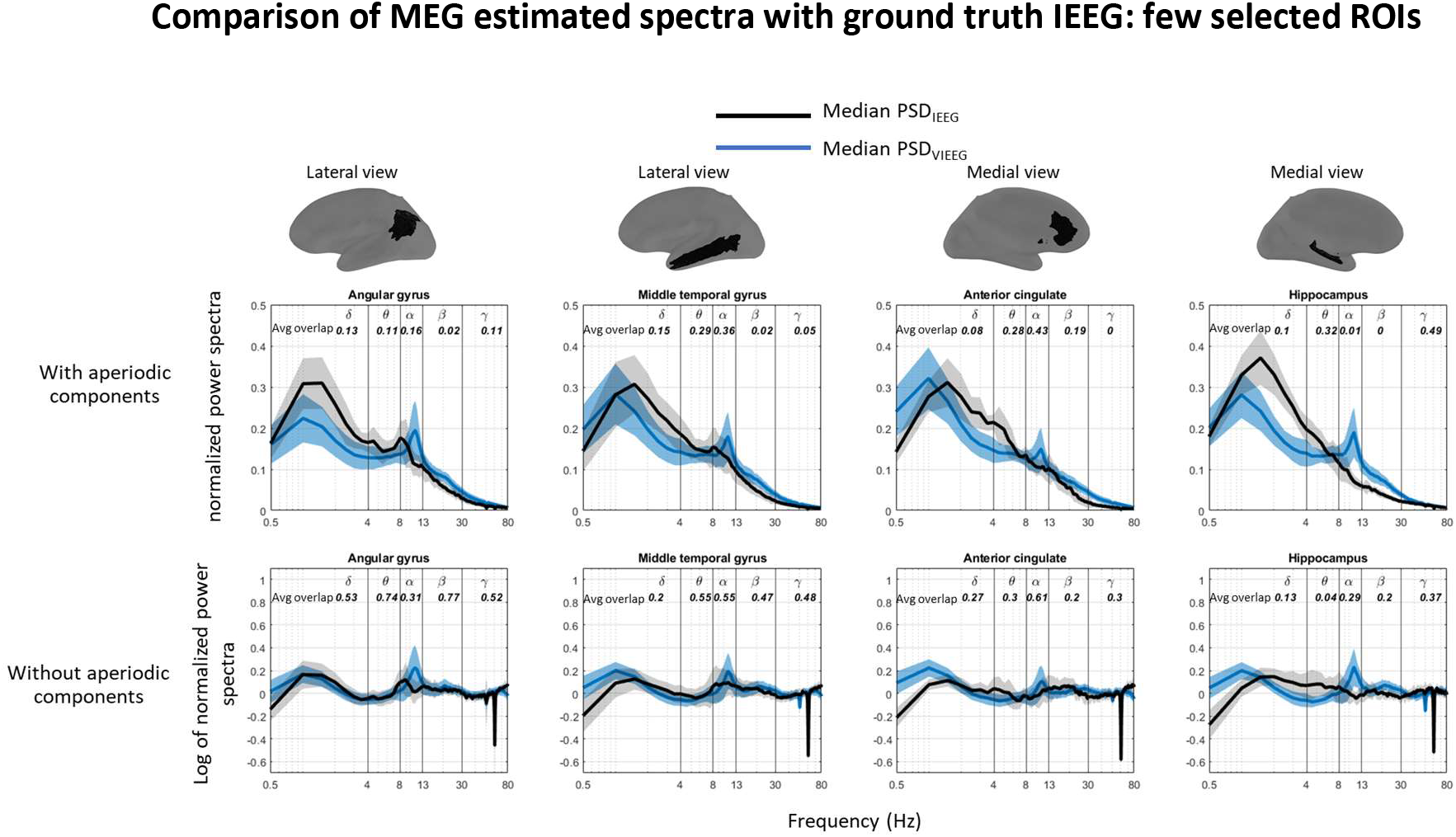
Comparison of periodic components of MEG estimated spectra with ground truth IEEG, with and without aperiodic components. Average overlaps between VIEEG and IEEG spectra across each spectral band are shown for a few selected ROIs. The value of *overlap* is calculated at each frequency bin and ranges from 0 to 1. For a ROI, if the median of PSD_VIEEG_ perfectly coincides with the median of PSD_IEEG_ at all frequency bins within a specific frequency band, the average *overlap* is 1.

For *angular gyrus,* the MEG estimated the spectra quite well compared to IEEG, in most frequency bands after removing the aperiodic components. The similarity between spectra was quantified by *average overlap* (*δ: 0.53, θ: 0.74, α: 0.31, β: 0.77, γ: 0.52*). Before removing the aperiodic components, the *average overlap* values in all frequency bands were clearly lower *δ: 0.13, θ: 0.11, α: 0.16, β: 0.02, γ: 0.11*.

In the *middle temporal gyrus*, the *average overlap* in all frequency bands after removing the aperiodic spectra are *δ: 0.2, θ: 0.55, α: 0.55, β: 0.47, γ: 0.48*. Here we chose another ROI for which MEG estimated VIEEG exhibited similar spectra to Gold Standard IEEG spectra, for all bands except delta. Before removing the aperiodic spectra, those values were much worse: *δ: 0.15, θ: 0.29, α: 0.36, β: 0.02, γ: 0.05*.

The example medial ROI *anterior cingulate* also showed improvement after removing the aperiodic components. The *average overlap* values in all frequency bands before and after removing aperiodic components are *δ: 0.08, θ: 28, α: 0.43, β: 0.19, γ: 0.0 and δ: 0.27, θ: 0.3, α: 0.61, β: 0.2, γ: 0.3*, respectively. For this medial structure, more difficult to localize in MEG because of its depth, *average overlap* values were good in the alpha band but lower in other bands (around 0.3), when compared to previous examples.

Finally, we showed an example of deep ROI, the *hippocampus.* MEG estimated the spectra in the *hippocampus* very poorly compared to IEEG. The *average overlap* values in all frequency bands before and after removing aperiodic components are *δ: 0.1, θ: 32, α: 0.01, β: 0, γ: 0.49 and δ: 0.13, θ: 04, α: 0.29, β: 0.2, γ: 0.37*, respectively. Although the spectral comparison improved after removing the aperiodic components, they remained inaccurate compared to other lateral superficial ROIs. It is worth mentioning that with MEG we estimated a clear peak in the alpha band in the hippocampus, whereas clearly IEEG data were exhibiting no alpha band peak.

The comparison between IEEG and VIEEG spectra after removing the aperiodic components for 38 ROIs is shown in Fig S3.

In Fig 5, we summarized the *average overlap* for all 38 ROIs before and after removing the aperiodic components. It shows for all frequency bands, the *average overlap* values improved after we removed the aperiodic components. If we compare the ROIs after removing the aperiodic spectra, we observe that the spectra in lateral regions are overall better estimated when compared to the medial ROIs. It was clearly the case for deeper regions like the hippocampus and amygdala, for which MEG estimated VIEEG spectra were not accurately recovering the actual IEEG spectra. On the other hand, the PSD in lateral temporal and parietal were localized accurately for all bands, as well as the medial posterior cingulate region. Especially, the medial ROIs in the theta band were very poorly estimated compared to other frequency bands. The delta band was very poorly estimated in occipital ROIs. This was also the case when we compared relative power in Fig 2, IEEG showed high activation in the delta band in the occipital regions which were not well estimated by MEG.

**Figure 5:**
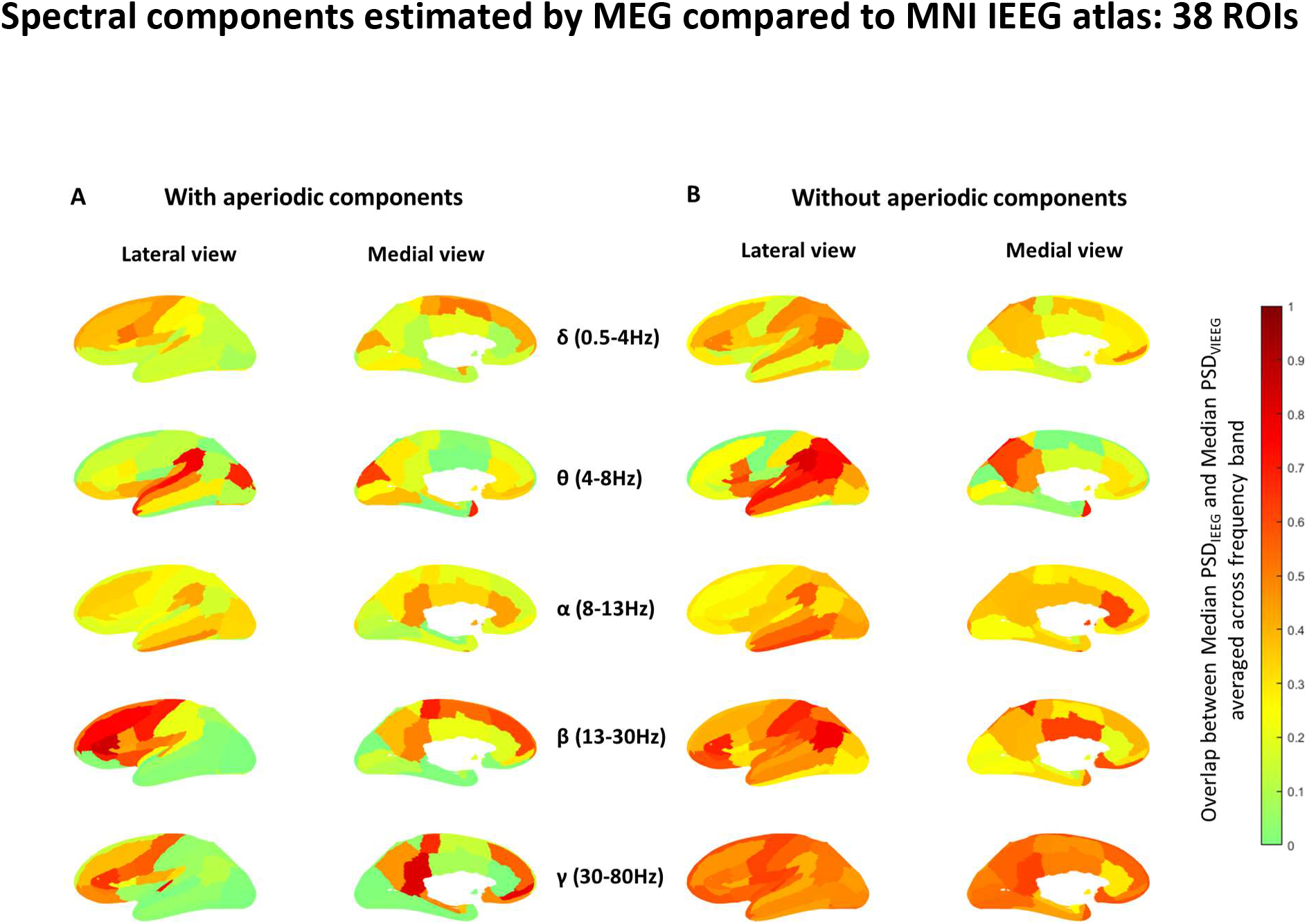
*Average overlap* between VIEEG and IEEG spectra across each spectral band for each of the 38 ROIs, (A) with and (B) without aperiodic components. The value of *overlap* is calculated at each frequency bin and ranges from 0 to 1. For a ROI, if the median of PSD_VIEEG_ perfectly coincides with the median of PSD_IEEG_ at all frequency bins within a specific frequency band, the average *overlap* is 1.

### 3.4 Comparison of VIEEG with IEEG in terms of oscillatory peaks

Oscillatory peaks in each band were estimated using the FOOOF algorithm, after removing the aperiodic component. When we compared MEG estimated spectra with the MNI IEEG atlas in terms of oscillatory peaks, we found that the probability histogram of peaks from all ROIs (Fig 6A) in IEEG has more variability in all frequency bands, whereas MEG estimated peaks are more narrowly concentrated within each frequency band, especially exhibiting high concentration in the alpha band. Also, the MEG estimated peaks in the theta band were much fewer than in IEEG. Fig 6B shows the probability histogram of peaks identified in IEEG, and VIEEG for an example ROI (hippocampus) for one subject. It also shows the value of the percentage difference of the number of channels exhibiting peaks in a specific frequency band, as a proportion of the total number of channels in that ROI *(Peak_estimated_SUBi_)* (Eq. 4). For instance, IEEG found peaks in the theta band, whereas no peak was found in VIEEG channels in this band. This was quantified in terms of the percentage difference of the number of channels exhibiting peak, *Peak_estimated_SUBi_* = −40% (underestimation) in the theta band. Thus, in the hippocampus, MEG clearly underestimated peaks in the delta, theta, and gamma band by 36%, 40%, and 31% respectively. On the other hand, we observed a large overestimation of channels exhibiting peaks in VIEEG in the alpha band by 83%. In the beta band, the estimation of peaks by VIEEG was comparable with those in IEEG *(Peak_estimated_SUBi_ = 0*). The probability histograms of peaks identified in IEEG and VIEEG for all 38 ROIs of 45 subjects are shown in Fig S4.

**Figure 6:**
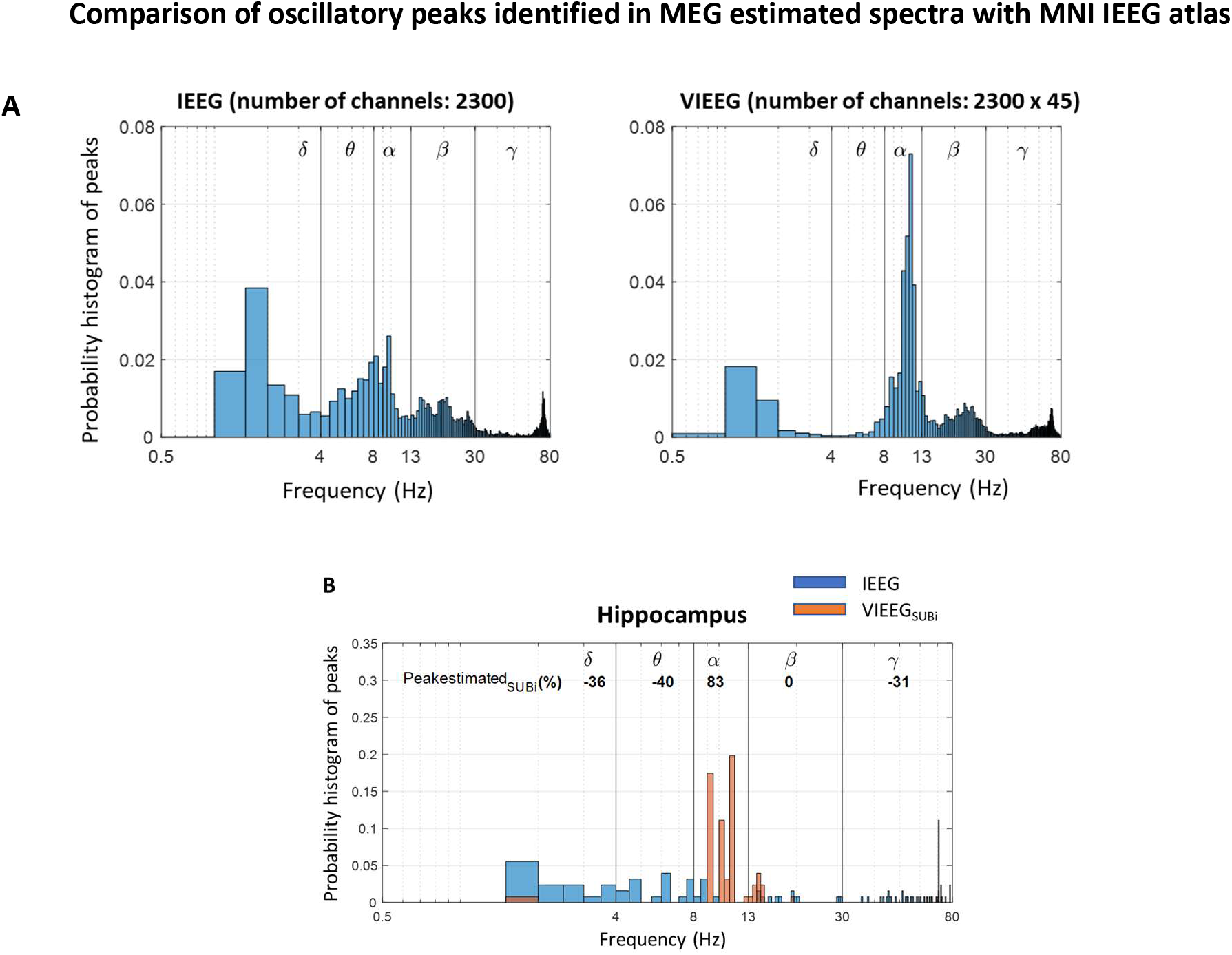
(A) Probability histogram of all identified peaks in all ROIs for IEEG and VIEEG using the FOOOF toolbox. (B) Percentage difference of the number of channels exhibiting spectral peaks in VIEEG compared to IEEG in each spectral band, as a proportion of the total number of channels in the ROI: illustration for the hippocampus ROI for one subject. The value of *Peak_estímated_SUBi_* ranges from −100% to 100%. The peaks are better estimated by VIEEG in comparison with IEEG, for a specific frequency band, if *Peak_estímated_SUBi_* is close to zero.

In Fig 7, we summarized the percentage difference of the number of channels exhibiting a peak in a specific frequency band for 45 subjects, by plotting the median of *Peak_estimated_SUBi_* (calculated for each subject) over 45 subjects and shown for 38 ROIs. Warmer colors indicate an overestimation and cooler colors indicate an underestimation of channels exhibiting peaks by MEG when compared to the MNI IEEG atlas. We observe that MEG overestimated peaks in the alpha band for most ROIs, especially higher in frontal (lateral and medial) ROIs (>40%) and deeper ROIs such as hippocampus and amygdala (~100% overestimation). The peaks in the delta band were well estimated in most ROIs except the occipital ROIs (like Fig 2 and Fig 5), and deep ROIs such as the hippocampus and posterior cingulate, where peaks were underestimated by MEG (<-35%). We observe an underestimation of peaks in beta and theta bands, in frontal and central ROIs (both lateral and medial) (<-35%). MEG moderately overestimated beta peaks in posterior regions and gamma peaks all over the brain regions. Those regions also showed higher relative power in MEG (Fig 2) compared to IEEG.

**Figure 7:**
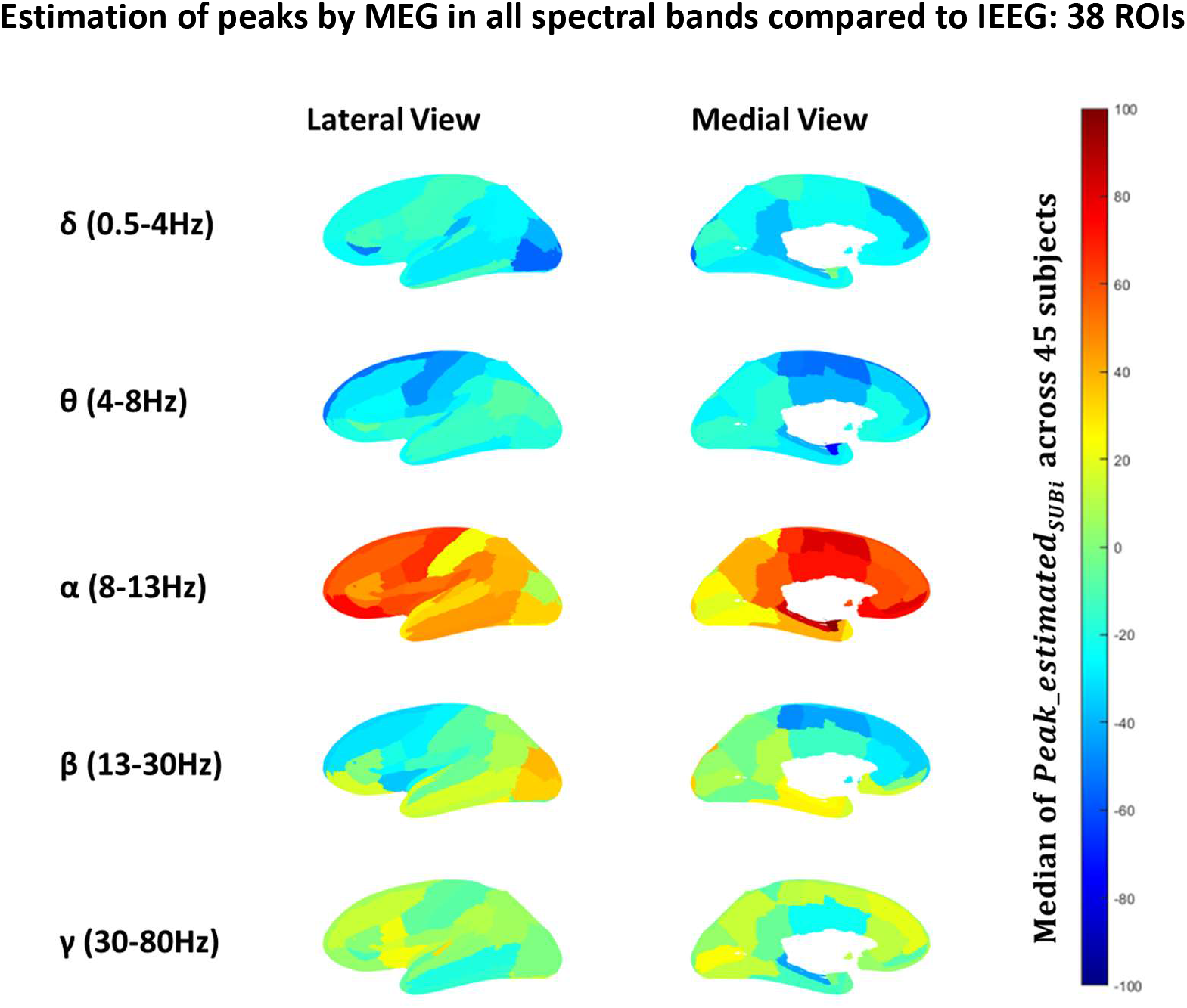
Median values of *Peak_estimated_SUBi_* over 45 subjects are plotted for 38 ROIs. *Peak_estimated_SUBi_* measures the percentage difference of the number of channels exhibiting spectral peaks in VIEEG compared to IEEG, as a proportion of the total number of channels in each ROI, for each spectral band and ROI, calculated for each subject *i*. The value of the *median Peak_estimated_SUBi_* over all subjects ranged from −100% to 100%. *Median (Peak_estimated_SUBi_)*=+100% indicates that all the channels from VIEEG in a ROI (N_ROI_) showed a peak in that frequency band, whereas no peak were identified in any of the IEEG channels in that ROI. We called it a 100% overestimation of oscillatory peaks by MEG estimated VIEEG in this ROI. On the contrary, a −100% estimation is obtained when all the IEEG channels in a ROI exhibit peaks, but no peak was identified in VIEEG in that ROI, resulting in a 100% underestimation. The peaks were better estimated by VIEEG for the ROIs if *median* (*Peak_estimated_SUBi_)* was close to zero. The values in the color bar are ranging from underestimation (−100%, negative values, cooler color) to overestimation (+100%, positive values, warmer color) of channels exhibiting peak by MEG estimated VIEEG.

It is also important to mention that all our results were reported in a common average montage. We also produced these results for bipolar montage and a similar pattern was found. Please see supplementary Fig S5 and Fig S6 comparing the results from the two montages. Although the spectral components recovered by both montages were very reproducible, bipolar montage was slightly better at estimating the oscillatory peaks in the alpha band, especially in the frontal regions.

### 3.5 IEEG and VIEEG amplitude

In Fig 8, the average amplitude across the IEEG and VIEEG channels in each ROI is plotted for the 38 ROIs, where each ROI amplitude was normalized by the average amplitude of all 38 ROIs. The mean amplitude across 38 ROIs was 28.4 *μ*V for IEEG, and 0.67 *μ*V for VIEEG, since underestimation of the amplitude after solving the MEG inverse problem was expected by the regularization procedure. Thus, we normalized IEEG and VIEEG amplitudes to be comparable. A strong positive correlation (Spearman’s *R* = 0.69, *p* < 0.001) was found for the amplitudes of 38 ROIs between IEEG and VIEEG. In Fig 8, we also plotted the difference of normalized amplitudes between IEEG and VIEEG for each ROI, called the *amplitude difference*. We represented the absolute value of *amplitude difference* in the bar plot and showed the signed amplitude difference on the inflated cortical surface. In the lateral occipital, lateral parietal, and lateral temporal regions, the VIEEG amplitudes were larger than the IEEG amplitudes. On the other hand, in the medial frontal regions and some deep regions such as the hippocampus and amygdala, the IEEG amplitudes were much higher than the MEG estimated VIEEG amplitudes.

**Figure 8:**
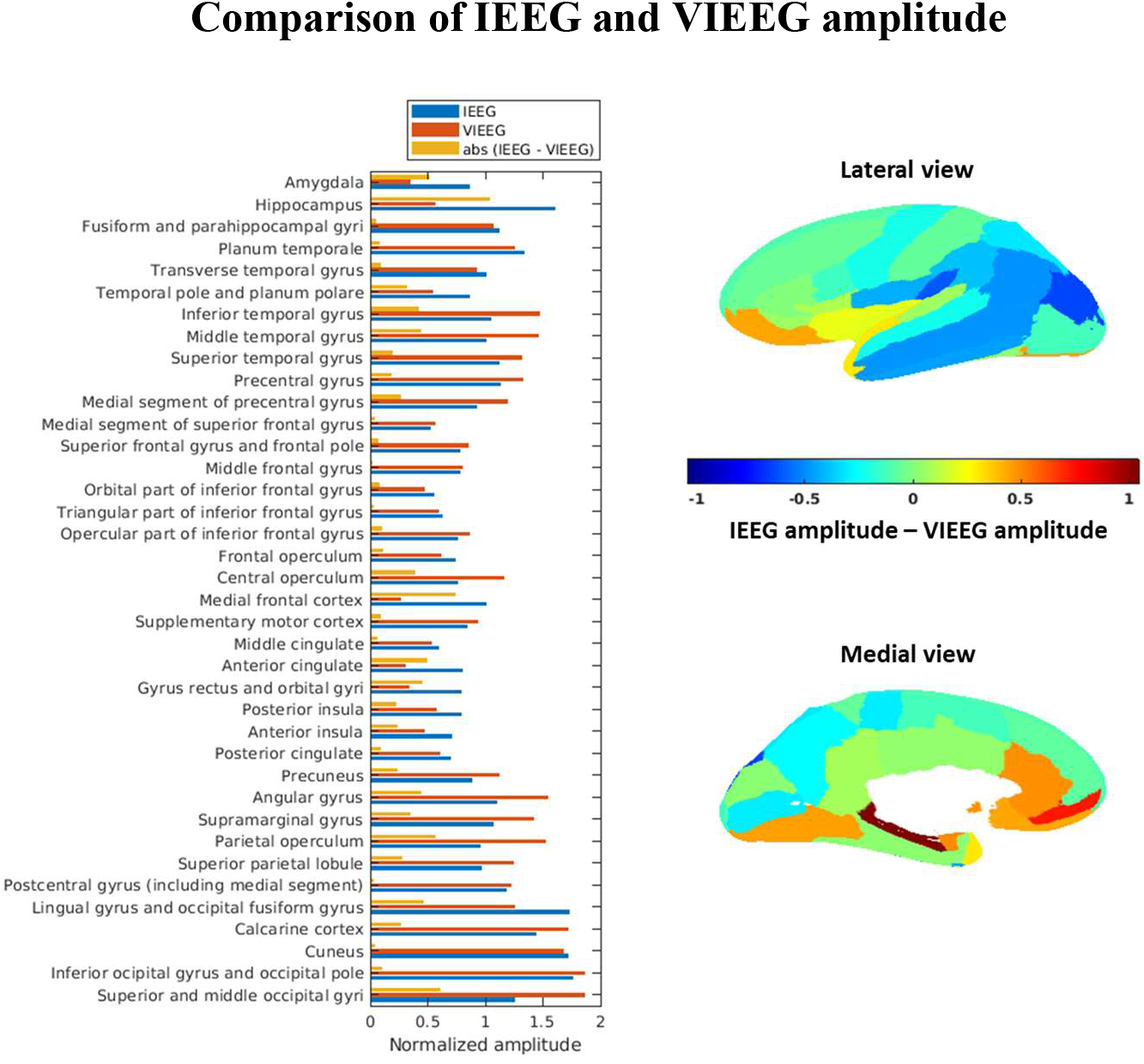
Average amplitude across the channels in each ROI, normalized by the average amplitude across all ROIs. The bar plot shows the absolute difference of normalized amplitude between IEEG and VIEEG. The right panel shows the difference of normalized amplitude between IEEG and VIEEG on the inflated cortical surface (lateral and medial views).

We calculated the correlation between the signed *amplitude difference* and the *average overlap* (calculated in Fig 5) for 38 ROIs in all frequency bands. It showed moderate negative correlation in alpha band (Spearman’s *R* = −0.33, *p* = 0.04) and beta band (Spearman’s *R* = −0.3, *p* = 0.08). A negative correlation indicates that for regions exhibiting higher VIEEG amplitude when compared to IEEG amplitudes, the *average overlap* between VIEEG and IEEG spectra were better. In delta and theta frequency bands, weak negative correlation was found (δ: Spearman’s *R* = −0.19, *p* = 0.25, θ: Spearman’s *R* = −0.22, *p* = 0.18). No correlation was found in gamma band (Spearman’s *R* = −0.03, *p* = 0.8).

## 4 Discussion

We aimed to assess the reliability of MEG source imaging of awake resting state oscillations by validating with the MNI IEEG atlas as ground truth (Frauscher et al., 2018). We compared MEG estimated VIEEG spectra from a healthy group of participants (Pellegrino et al., 2022) with the MNI IEEG atlas of healthy brain awake activity, in terms of (i) oscillatory components of the spectra, (ii) oscillatory peaks, and (iii) relative power. This is the first study using an IEEG atlas of healthy awake activity to validate quantitatively the accuracy of MEG source imaging of resting state activity. In this context, we carefully investigated the performance of our source imaging technique, the wavelet based Maximum Entropy on the Mean (wMEM) (Aydin et al., 2020; Lina et al., 2012; Pellegrino et al., 2016). A quantitative comparison between IEEG and VIEEG spectra showed that the VIEEG spectra were closer to the IEEG spectra after the aperiodic components were removed from the spectra (Fig 5). The estimation of the VIEEG spectra was overall more accurate in the lateral regions compared to the medial regions (Fig 5B). Especially better estimation was found in the regions exhibiting higher VIEEG amplitude compared to IEEG (Fig 8), such as the lateral parietal, lateral temporal, and some lateral occipital regions (Fig 5B). Importantly, we found that the estimation of VIEEG resting state spectra was particularly inaccurate in the deep regions such as the hippocampus and amygdala, for most frequency bands. Our study also found that MEG estimated spectra were dominated by oscillations in the alpha band, especially in anterior and deeper regions, unlike the actual in situ measurements from the MNI IEEG atlas (Fig 4, Fig 7, Fig S3). This observation is consistent with the finding of dominance in alpha oscillations reported in previous studies (Capilla et al., 2022; Keitel & Gross, 2016; Mahjoory et al., 2020). In our study, the MNI IEEG atlas as ground truth enabled us to quantify the extent of overestimation or underestimation of alpha dominance in MEG estimated spectra. A quantitative comparison of oscillatory peaks showed that MEG overestimated peaks in the alpha band in most brain regions, especially in the frontal and deep regions (Fig 7). In the delta, theta, and gamma bands, the peaks in the deep regions were underestimated, whereas they were more accurately estimated in lateral cortical regions. In terms of relative power, the distribution of MEG relative power reported in our study was consistent with the previous MEG studies in different frequency bands (Fig 2, Fig S1, Fig S2) (Hillebrand et al., 2012; Mahjoory et al., 2020; Mellem et al., 2017; Niso et al., 2016; Niso et al., 2019). However, when compared to the MNI IEEG atlas (Fig 2), important differences in average relative power were observed in the anterior regions for the alpha band, in the posterior regions for the delta, beta and gamma bands, and in deep regions (such as hippocampus and amygdala) for all bands. Especially in the theta band, MEG largely underestimated relative power compared to IEEG in all brain regions.

We also calculated the relative power for MEG using another source imaging method, depth weighted MNE (Fig 2, Fig S1, and Fig S2), to determine if the findings of our study were not mainly driven by our source imaging method, wMEM. Indeed, results were overall similar when depth weighted MNE was applied. Investigating more carefully the comparison between MNE and wMEM, VIEEG estimated using wMEM exhibited slightly better distribution (i.e., closer to IEEG) in relative power, when compared to MNE, especially in the theta band and gamma bands, for which MNE was respectively underestimating and overestimating the relative power.

### 4.1 Removing the aperiodic component improves the spectral comparison between VIEEG and IEEG

Electrophysiological power spectra are composed of periodic components, typically characterized by spectral peaks in canonical frequency bands and aperiodic components, also known as 1/f-like or arrhythmic components (He, 2014). While analyzing electrophysiological power spectra of neuronal oscillations, the separation of periodic and aperiodic components allows a better estimation of the periodic component (Donoghue et al., 2021; Wen & Liu, 2016). We applied FOOOF (Donoghue et al., 2020) to separate the periodic and aperiodic components of the spectra from IEEG and VIEEG. We quantitatively compared the IEEG and VIEEG spectra before and after removing the aperiodic components. The spectra estimated by MEG became more comparable to IEEG after the aperiodic components were removed (Fig 5). This is reflected by the metric *average overlap,* which quantifies the overlap between IEEG and VIEEG spectra. In Fig 5A, the *average overlap* values of the spectra, which include the aperiodic components, were very low for all frequency bands, with most ROIs exhibiting an *average overlap* < 0.2. The *average overlap* values of the ROIs were much improved after removing the aperiodic components (Fig 5B) compared to Fig 5A. This indicates that the discrepancies between IEEG and VIEEG spectra were mostly driven by the variations in the aperiodic components between the two modalities. The aperiodic component of the spectrum might be generated by spatial interactions among neuronal populations (Aguilar-Velázquez & Guzmán-Vargas, 2019). The IEEG records brain activity locally, with a spatial sensitivity of less than 1cm (von Ellenrieder et al., 2021), and would therefore not pick-up all the spatial interactions, leading to a different aperiodic component from the one recorded by the more spatially spread sensitivity of scalp EEG or MEG. Moreover, in a recent study based on computational modeling using the virtual brain project, we found that anatomical and forward model properties of EEG and MEG resulted in different aperiodic components between EEG and MEG (Bénar et al., 2019).

The differences in the aperiodic components between MEG and IEEG could also result from the differences in the inherent mechanism of generation of the signals in those two modalities. The local bioelectrical environment contributes to the generation of the local field potential of IEEG (Bédard & Destexhe, 2009), whereas MEG is mainly contributed by the activation of pyramidal sources.

### 4.2 MEG spectra were better estimated in the lateral regions compared to the medial regions

We observed that MEG estimated spectra were better estimated in lateral regions compared to the medial regions in most frequency bands (Fig 5B). The lateral parietal and lateral temporal regions in most frequency bands showed *average overlap* values greater than 0.5. In contrast, for most of the medial ROIs, the *average overlap* values were less than 0.5 in all frequency bands (except gamma). We found a negative correlation between the signed difference of IEEG and VIEEG amplitude (Fig 8) and the *average overlap* values (Fig 5B) in all frequency bands except gamma. These negative correlations were moderate in the alpha and beta bands, and weak in the delta and theta bands. Such negative correlations indicate that *average overlap* values were better in regions having VIEEG amplitudes greater than IEEG amplitudes, which means MEG could estimate the spectra from these ROIs more accurately when underlying signals were of larger amplitudes for those regions. Example regions include the lateral parietal, lateral temporal, and some lateral occipital regions, which exhibited VIEEG amplitudes much higher than IEEG amplitudes (Fig 8) and also resulted in higher *average overlap* (Fig 5B). Overall, an important finding was that deep regions were not well estimated by MEG, and we found similar results for wMEM and MNE (results not shown). For instance, the *average overlap* values were less than 0.3 in the hippocampus for all frequency bands except gamma, and less than 0.25 in the amygdala for all frequency bands. The reason for such poor estimation could be the large underestimation of the VIEEG amplitude compared to IEEG (Fig 8). Similarly, in the lingual gyrus and occipital fusiform gyrus, the VIEEG amplitude was largely underestimated compared to IEEG (Fig 8). The *average overlap* in this ROI was less than 0.25 in all frequency bands except gamma.

### 4.3 Dominance of alpha oscillations in MEG

Oscillatory peaks in the alpha band were largely overestimated by MEG, especially in frontal regions *(Median (Peak_estimated_SUBi_)* = ~45-84%) and deep regions such as the hippocampus *(Median (Peak_estimated_SUBi_) =* 86%) and amygdala *(Median (Peak_estimated_SUBi_)* = 100%). We found widespread alpha oscillations in all brain regions (Fig 7, Fig S3, Fig S4). A quantitative comparison between the spectra from MEG estimated VIEEG and the MNI IEEG atlas (Fig 5B) showed, in most of the lateral frontal regions and medial regions, an *average overlap* in the alpha band of less than 0.5. A similar dominance of alpha oscillations in MEG was also reported in previous MEG resting state studies (Capilla et al., 2022; Keitel & Gross, 2016; Mahjoory et al., 2020). Using eyes open MEG data, Capilla et al. (2022) reported the dominance of alpha oscillations in all posterior regions. Alpha is also of much higher amplitude with eyes closed than with eyes open. Thus, with eyes closed data, the dominance of alpha oscillations is expected to be more widespread and could explain why we found an overestimation of the alpha peak identified in MEG in most frontal brain regions. On the other hand, the large predominance of alpha oscillations found in deeper brain regions, that were also showing weak amplitudes, could be explained by spatial leakage from cortical signals getting localized with very little amplitude in deep regions. Nunez et al. (2001) and Srinivasan et al. (2006) also reported alpha dominance in brain regions with scalp recordings, including frontal regions. It is quite evident from intracranial EEG that alpha is not as prominent and widespread as seen from scalp recordings, especially not in the frontal regions (Groppe et al., 2013; Penfield & Jasper, 1954).

With electrocorticography (ECoG) recordings in patients with epilepsy, Groppe et al. (2013) reported that most dominant oscillations tended to be around ~7 Hz (in the theta range), not in the alpha range (8-13Hz) typically reported in scalp recordings. This is also evident from the peak histogram of IEEG and MEG (Fig 6A), IEEG tends to have the highest number of peaks around ~7Hz, within theta and alpha bands, whereas MEG peaks were around ~10-12Hz, in much higher proportions. Compared to IEEG, MEG underestimated theta and overestimated alpha, which was also evident in the relative power calculated in Fig 2. The reason EEG/MEG sees higher alpha oscillations might be due to a phase synchronization over larger spatial extents than other bands (Groppe et al., 2013). The IEEG having a very local sensitivity profile, would pick up the activity from the alpha band, but also from other bands with low spatial phase synchronization. MEG, on the other hand, having a more extended spatial profile, would pick up the generators of synchronous alpha activity interfering constructively, but generators of theta activity (or other poorly synchronized bands) would partially cancel out, leading to a dominant alpha rhythm.

To tackle the dominance of alpha oscillations, a few previous studies normalized each ROI spectrum by considering the average spectra of all other brain regions. Such normalization gives a measure of the characteristic features of each ROI spectrum compared to other brain regions, resulting in a less widespread influence of alpha oscillations in all brain regions (Capilla et al., 2022; Keitel & Gross, 2016). We did not incorporate such normalization in this study, as we aimed to compare the MEG estimated spectra for each ROI with the MNI IEEG atlas, not with other ROIs. We also investigated IEEG and VIEEG data in a bipolar montage (Fig S5 and Fig S6). The spectral components estimated by MEG compared to the MNI IEEG atlas were very similar for bipolar and common-average montages (Fig S5). However, the peaks estimated by MEG were less dominant in the frontal regions in the alpha band in the bipolar montage, when compared to a common-average reference montage (Fig S6).

### 4.4 Differences in signal relative power

A qualitative comparison of relative power between MEG estimated spectra and the MNI IEEG atlas (Fig 2, Fig S1) showed that in general, both modalities have similar brain distributions, such as strong delta and theta power in frontal regions, alpha power in posterior regions, beta power in motor and frontal regions, and gamma power in frontal areas. However, when compared to the MNI IEEG atlas, the MEG estimated relative power was spatially much more smoothly distributed. For instance, the IEEG atlas showed low power in the hippocampus in theta, alpha, and beta bands, in contrast to strong power in its neighboring region the para-hippocampal gyrus. In MEG, due to source leakage, such separation was not possible, resulting in similar distributions in the parahippocampal gyrus, the hippocampus, and the neighboring regions for all frequency bands (Fig 2, Fig S1). The contrast between strong and weak relative power was reflected in MEG only where an extended area in IEEG exhibited a similar contrast. For instance, the frontal regions showed weak alpha power compared to the posterior regions in IEEG, a pattern that was also found in VIEEG (Fig S1). Similarly, posterior regions in the theta band and the orbito-frontal region in the beta band showed weak power compared to other brain regions within the specific band in IEEG, which were also reflected in VIEEG (Fig S1). Due to the source leakage in MSI, subtle changes in relative power in brain regions (as seen in the MNI IEEG atlas) were not accurately retrieved using MEG (Fig 2). This was particularly the case for deep regions localized with small amplitude in MEG. Similar mislocalization patterns were found with wMEM and MNE.

Our MEG relative power maps were quite consistent with previous MEG studies (Niso et al., 2016; Niso et al., 2019). We also compared the MEG relative power on the cortical surface and MEG estimated VIEEG relative power in the intracranial space in Fig S2 (described in the supplementary material). Fig S2 confirms that the conversion from the MEG source map to intracranial space (VIEEG) did not add any discrepancy, and the relative power in the virtual intracranial space (VIEEG) was concordant with the MEG relative power on the cortical surface. We also included source imaging results from depth weighted MNE in Fig 2, Fig S1, and Fig S2, to show that the findings in our study were not driven by our source imaging method, wMEM. We get similar findings with depth weighted MNE (as discussed in the supplementary material). It is worth mentioning that our method consisting in converting MEG sources into virtual IEEG potentials (Abdallah et al., 2022; Grova et al., 2016) offers a solid quantification approach to compare MEG sources estimated using different source imaging techniques with actual IEEG in situ recordings, carefully taking into account spatial sampling of IEEG data.

### 4.5 wMEM for resting state localization and comparison with MNE

We implemented and validated an adapted version of wavelet MEM to solve the inverse problem in the context of resting state source imaging. wMEM is a MEM framework specifically designed to localize oscillatory brain patterns in the context of EEG/MEG signals utilizing discrete wavelet transformation (Daubechies wavelets). Taking advantage of the MEM specific prior model (Chowdhury et al., 2013), wMEM can accurately localize the oscillatory patterns together with their spatial extent. This ability was validated for localizing oscillatory patterns at seizure onset (Pellegrino et al., 2016), interictal bursts of high frequency oscillations (Avigdor et al., 2021; von Ellenrieder et al., 2016) as well as for localizing MEG resting state fluctuations in the alpha band (Aydin et al., 2020). We further adapted wMEM to localize wide band oscillations in resting state EEG/MEG data. (i) Spatial prior model: Our main adaptation of the wMEM spatial prior model consisted in considering one stable whole brain parcellation of the cortical surface, following the strategy proposed for cMEM (Chowdhury et al., 2013), whereas in our previous wMEM implementation the parcellation was varying for time frequency samples. To do so, we proposed data-driven whole brain parcellation informed by the MSP method (Mattout et al., 2005), a projection technique allowing to estimate the probability of every source to contribute to the data, before region growing around local MSP peaks. The main adaptation of our current implementation is that the MSP projector is applied on all wavelet coefficients of Daubechies timefrequency representation of our data, instead of using signals in the time domain (see detailed in the Appendix). (ii) Initialization of the probability of being active for each parcel: Following the parcellation, we then initialized the probability of each parcel of being active or not, using normalized energy calculated for each time frequency sample. (iii) Selection of baseline for resting state localization: There is no ideal baseline definition when localizing ongoing resting state data. We proposed to compute the sensor level noise covariance matrix from the ongoing resting state data. To do this, we generated a quasi-synthetic baseline from the signal of interest by randomly modifying the Fourier phase at each frequency. We also adopted a sliding window approach to generate the baseline for a more accurate estimation of the noise covariance matrix for each time frequency sample along the time scale. More details of these adaptations are described in the Appendix.

Before applying this new implementation of wMEM on resting state MEG data, we validated it within a controlled environment with realistic simulation of epileptic spikes and oscillations on realistic MEG background, as previously proposed in Chowdhury et al. (2013) and Lina et al. (2012). We observed that this new wMEM demonstrated improvement in localizing the underlying spatial extent of the generators when compared to the previous wMEM implementation (Lina et al., 2012) and depth weighted MNE (results not shown), therefore justifying our rationale for incorporating the adaptations when localizing resting state data.

In the present study, we also compared wMEM and depth weighted MNE when localizing resting state MEG data. The comparison in terms of relative power (Fig S2) showed similar maps in all frequency bands except gamma. However, when compared to actual IEEG power (Fig 2), VIEEG estimated using wMEM exhibited better distribution in relative power, when compared to MNE, especially in theta and gamma bands. VIEEG computed from MNE sources showed large underestimation in the theta band and overestimation in the gamma band, whereas VIEEG power estimated from wMEM sources recovered power in the theta and gamma bands more accurately. Additionally, the relative power in the delta band was slightly better estimated using wMEM compared to depth weighted MNE. The reason wMEM localized brain activity in the lower (delta and theta) and higher (gamma) frequency range more accurately might be explained by the sparse/optimal data representation of oscillatory components obtained when using Daubechies discrete wavelets (Lina et al., 2012), before solving the inverse problem.

It is also important to mention that, unlike wMEM, depth weighted MNE is designed to compensate for the bias of high sensitivity of the forward model towards superficial sources. Nevertheless, our overall results using both methods were similar. Currently, we are implementing depth weighting in wMEM, following a strategy proposed in Cai et al. (2022) for MEM implemented for the reconstruction of functional Near-InfraRed Spectroscopy data. In addition, we are including deep subcortical structures (such as the hippocampus) in the MEM model. We expect that incorporating depth weighted wMEM with more accurate models of subcortical structures (Attal & Schwartz, 2013) could improve the localization of resting state oscillation in deep regions. This investigation was out of the scope of the present study.

### 4.6 Limitations

One limitation of this study is that the normative MNI IEEG atlas was collected from patients with epilepsy, although by including only the IEEG electrodes implanted in the brain regions which turned out to be healthy. This limitation cannot be overcome, as IEEG data are never collected from healthy subjects. Nevertheless, such normative IEEG atlas data are so far the best ground truth and provide us with a unique opportunity to validate non-invasive source imaging techniques. Another limitation is the heterogeneity of IEEG channels in different ROIs. The sampling of IEEG channels was higher in lateral temporal, parietal, and frontal regions compared to medial and occipital regions. This might have biased our results, but not severely. We found a mild to moderate positive correlation between the number of channels in a ROI and the *average overlap* value (calculated in Fig 5B) in delta and beta bands (results not shown). We also combined channels in both hemispheres to maximize the sampling and the coverage of the brain. Thus, any effect of hemispheric asymmetry on oscillatory characteristics was lost. Also, the MEG data and IEEG data in this study were not simultaneously recorded. Simultaneous IEEG-MEG recordings would give us more opportunities to validate region specific spectral components (De Stefano et al., 2022; Pizzo et al., 2019), which we plan to do in the future. However simultaneous IEEG-MEG recordings have limited spatial sampling, whereas we could study the whole brain with the normative MNI IEEG atlas.

## 5 Conclusion

We aimed to address the reliability of MEG source imaging by validating source imaging results with the MNI IEEG atlas as ground truth. We quantitatively estimated the concordance of MEG estimated spectral components with the MNI IEEG atlas and identified the brain regions for which MEG estimated spectra are reliable and some regions for which we should be cautious while interpreting MEG results. We found widespread leakage of alpha oscillations in MEG estimated spectra in frontal and deep brain regions, which was present both before and after the removal of aperiodic components. In the future, we are planning to investigate these issues on simultaneous MEG-IEEG data and validate MEG source imaging of spectral components at the single-subject level.

## Acknowledgment

This work was supported by Natural Sciences and Engineering Research Council of Canada (NSERC) Discovery grant, grant from Canadian Institutes of Health Research (CIHR) (PJT-159948 and FDN 143208), and the Fonds de recherche du Québec—Nature et technologies (FRQNT) Research team grant.

## Appendix

The maximum entropy on the mean (MEM) is a Bayesian inference technique that regularizes the inverse problem using prior information. This prior relies on the notion of functional parcellation of brain activity over the cortical surface, and hidden state variables describing each parcel being active or inactive. A data driven parcellization (DDP) based on Multivariate Source Prelocalization (MSP) method (Mattout et al., 2005) was used to guide the parcellation of the cortical surface into non-overlapping and functionally homogeneous parcels (Lapalme et al., 2006). For each parcel, the prior is then defined as a mixture of Gaussians, each Gaussian of each parcel will be related to a state as active and inactive controlled by a hidden state variable. In the context of resting state EEG/MEG, we adapted wMEM by incorporating a few changes in the prior model and initialization of the parcels.

### Spatial prior model

Parcellation of the whole cortical surface was obtained using a data driven approach, based on the Multivariate Source Pre-localization (MSP) method (Mattout et al., 2005), a projection technique allowing to estimate the probability of every source contributing to the data. The MEG data (M) were first normalized (across sensors) and then wavelet transformed 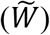. In the present implementation of wMEM, time expansion was thus substituted with a time-scale representation. The contribution of each source to the data (called the MSP score) is obtained using the normalized MEG data 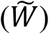 and the normalized lead field matrix 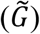, where the normalization was performed by the norm of each column. 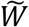 is the sensor space data in the wavelet domain (dimension: *number of sensors × discrete wavelet time-frequency indices),* whereas the normalized lead field 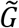 is of dimension *number of sensors × n; n* is the total number of sources. The MSP score for each source (ai), *i*= 1,…. *n*, is calculated in the following steps:

The normalized lead field matrix 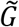 is decomposed into *d* mutually orthogonal eigenvectors *ui* using singular value decomposition (SVD). We selected a subspace, U_s_ = [u_1_, u_2_,… u_s_], by projecting these orthogonalized projectors onto the normalized data 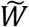 as 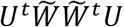 and taking the diagonal of 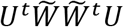 that captures 95% of the variability.

In the data subspace spanned by U_s_, the data that can be explained within this subspace is calculated as:

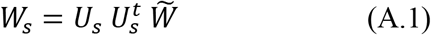

The projector in this subspace is defined by

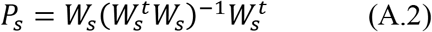

Finally, the MSP score between 0 and 1 for each source *i* is then calculated by the norm of the projection of its associated lead field,

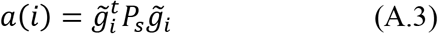

where 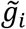 is the *i^th^* column of 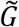. Parcels are then constructed using a region-growing algorithm, selecting sources according to decreasing MSP scores. In this version of wMEM, assuming a stable parcellation of the cortex for all the time-frequency samples for resting state data, we followed the strategy proposed for cMEM (Chowdhury et al., 2013), ensuring that the same underlying parcellation was considered when localizing all the time-frequency samples (dimension of 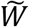 as *number of sensors × number of discrete time frequency boxes).* In our previous implementation (Lina et al., 2012), a specific parcellation was computed for each time-frequency box to localize.

### Initialization of the parcels

The probability *α_k_* for each parcel *k* to be active was then initialized as the amount of ‘normalized energy’ in the parcel, for each time-frequency sample. Given the minimum norm estimated energy of the sources, for a specific time-frequency sample (a column in 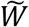) 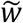,

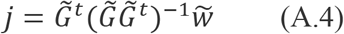

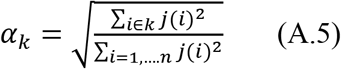

Although the parcels were identical across time-frequency samples, this quantity initializing the probability of each parcel to be active changed with time and frequency.

### Selection of baseline for resting state localization

A baseline is needed to complete the initialization of the prior, to define the variance of the active state and inactive state, in comparison to the noise variance at the sensor level. This is an important feature that will allow switching off parcels in the model when they are not active. The idea of selecting a baseline is to choose a segment of data with an amplitude significantly lower than the signal of interest. However, the selection of such segments in resting state data is not straightforward. In Aydin et al. (2020), the baseline was defined as a two-second segment exhibiting low amplitude in the alpha band, since we were investigating amplitude envelope correlation in the alpha band, as a connectivity metric. This approach worked reasonably when localizing in a specific and narrow frequency band, however, becomes inappropriate when localizing in a wide frequency band. Here, we propose to generate a quasi-synthetic baseline from a segment of the signal of interest. The baseline was obtained by randomly modifying the Fourier phase at each frequency, for all the sensors (originally proposed by Prichard & Theiler 1994). This baseline preserves the coherence between the sensors and the power spectrum of the signals while destroying only the temporal coherence. To consider this new baseline, we adopted a sliding window approach to calculate the baseline. For each window with one second duration along the sixty second resting state MEG data, a quasi synthetic “shuffled” baseline was thus generated. To solve the inverse problem for each time frequency box, we selected the corresponding one second “shuffled” baseline along the time scale. We adopted this sliding window approach considering that the selection of baseline is an important aspect of the initialization of the prior, allowing the parcels to be active or not, and thus would be more reasonable to use the baseline which is temporally associated with the time frequency sample.

## References

Abdallah, C., Hedrich, T., Koupparis, A., Afnan, J., Hall, J. A., Gotman, J., Dubeau, F., von Ellenrieder, N., Frauscher, B., & Kobayashi, E. (2022). Clinical Yield of Electromagnetic Source Imaging and Hemodynamic Responses in Epilepsy: Validation With Intracerebral Data. Neurology.

Aguilar-Velázquez, D., & Guzmán-Vargas, L. (2019). Critical synchronization and 1/f noise in inhibitory/excitatory rich-club neural networks. Scientific reports, 9(1), 1–13.

Amblard, C., Lapalme, E., & Lina, J.-M. (2004). Biomagnetic source detection by maximum entropy and graphical models. IEEE transactions on biomedical engineering, 51(3), 427–442.

Attal, Y., & Schwartz, D. (2013). Assessment of subcortical source localization using deep brain activity imaging model with minimum norm operators: a MEG study. PloS one, 8(3), e59856.

Avigdor, T., Abdallah, C., von Ellenrieder, N., Hedrich, T., Rubino, A., Russo, G. L., Bernhardt, B., Nobili, L., Grova, C., & Frauscher, B. (2021). Fast oscillations> 40 Hz localize the epileptogenic zone: An electrical source imaging study using high-density electroencephalography. Clinical Neurophysiology, 132(2), 568–580.

Aydin, Ü., Pellegrino, G., Ali, O. B. K. b., Abdallah, C., Dubeau, F., Lina, J.-M., Kobayashi, E., & Grova, C. (2020). Magnetoencephalography resting state connectivity patterns as indicatives of surgical outcome in epilepsy patients. Journal of neural engineering, 17(3), 035007.

Badier, J.-M., Dubarry, A., Gavaret, M., Chen, S., Trébuchon, A., Marquis, P., Régis, J., Bartolomei, F., Bénar, C.-G., & Carron, R. (2017). Technical solutions for simultaneous MEG and SEEG recordings: towards routine clinical use. Physiological Measurement, 38(10), N118.

Becker, H., Albera, L., Comon, P., Gribonval, R., Wendling, F., & Merlet, I. (2015). Brain-source imaging: From sparse to tensor models. IEEE Signal Processing Magazine, 32(6), 100–112.

Bédard, C., & Destexhe, A. (2009). Macroscopic models of local field potentials and the apparent 1/f noise in brain activity. Biophysical journal, 96(7), 2589–2603.

Bénar, C. G., Grova, C., Jirsa, V. K., & Lina, J.-M. (2019). Differences in MEG and EEG power-law scaling explained by a coupling between spatial coherence and frequency: a simulation study. Journal of computational neuroscience, 47(1), 31–41.

Bódizs, R., Szalárdy, O., Horváth, C., Ujma, P. P., Gombos, F., Simor, P., Pótári, A., Zeising, M., Steiger, A., & Dresler, M. (2021). A set of composite, non-redundant EEG measures of NREM sleep based on the power law scaling of the Fourier spectrum. Scientific reports, 11(1), 1–18.

Brookes, M. J., Woolrich, M., Luckhoo, H., Price, D., Hale, J. R., Stephenson, M. C., Barnes, G. R., Smith, S. M., & Morris, P. G. (2011). Investigating the electrophysiological basis of resting state networks using magnetoencephalography. Proceedings of the National Academy of Sciences, 108(40), 16783–16788.

Buzsáki, G., Logothetis, N., & Singer, W. (2013). Scaling brain size, keeping timing: evolutionary preservation of brain rhythms. Neuron, 80(3), 751–764.

Cai, Z., Machado, A., Chowdhury, R. A., Spilkin, A., Vincent, T., Aydin, Ü., Pellegrino, G., Lina, J.-M., & Grova, C. (2022). Diffuse optical reconstructions of functional near infrared spectroscopy data using maximum entropy on the mean. Scientific reports, 12(1), 1–18.

Capilla, A., Arana, L., García-Huéscar, M., Melcón, M., Gross, J., & Campo, P. (2022). The natural frequencies of the resting human brain: an MEG-based atlas. NeuroImage, 119373.

Chowdhury, R., Merlet, I., Birot, G., Kobayashi, E., Nica, A., Biraben, A., Wendling, F., Lina, J.-M., Albera, L., & Grova, C. (2016). Complex patterns of spatially extended generators of epileptic activity: Comparison of source localization methods cMEM and 4-ExSo-MUSIC on high resolution EEG and MEG data. NeuroImage, 143, 175–195.

Chowdhury, R. A., Lina, J. M., Kobayashi, E., & Grova, C. (2013). MEG source localization of spatially extended generators of epileptic activity: comparing entropic and hierarchical bayesian approaches. PloS one, 8(2), e55969.

Collins, D. L., Neelin, P., Peters, T. M., & Evans, A. C. (1994). Automatic 3D intersubject registration of MR volumetric data in standardized Talairach space. Journal of computer assisted tomography, 18(2), 192–205.

Cosandier-Rimélé, D., Badier, J.-M., Chauvel, P., & Wendling, F. (2007). A physiologically plausible spatiotemporal model for EEG signals recorded with intracerebral electrodes in human partial epilepsy. IEEE transactions on biomedical engineering, 54(3), 380–388.

Dalal, S., Jerbi, K., Bertrand, O., Adam, C., Ducorps, A., Schwartz, D., Martinerie, J., & Lachaux, J.-P. (2013). Simultaneous MEG-intracranial EEG: new insights into the ability of MEG to capture oscillatory modulations in the neocortex and the hippocampus. Epilepsy & Behavior, 10.1016/j.yebeh.2013.1003.1012.

Dale, A. M., Fischl, B., & Sereno, M. I. (1999). Cortical surface-based analysis: I. Segmentation and surface reconstruction. NeuroImage, 9(2), 179–194.

Darvas, F., Pantazis, D., Kucukaltun-Yildirim, E., & Leahy, R. (2004). Mapping human brain function with MEG and EEG: methods and validation. NeuroImage, 23, S289–S299.

De Stefano, P., Carboni, M., Marquis, R., Spinelli, L., Seeck, M., & Vulliemoz, S. (2022). Increased delta power as a scalp marker of epileptic activity: a simultaneous scalp and intracranial electroencephalography study. European journal of neurology, 29(1), 26–35.

Donoghue, T., Haller, M., Peterson, E. J., Varma, P., Sebastian, P., Gao, R., Noto, T., Lara, A. H., Wallis, J. D., & Knight, R. T. (2020). Parameterizing neural power spectra into periodic and aperiodic components. Nature neuroscience, 23(12), 1655–1665.

Donoghue, T., Schaworonkow, N., & Voytek, B. (2021). Methodological considerations for studying neural oscillations. European journal of neuroscience.

Dubarry, A.-S., Badier, J.-M., Trébuchon-Da Fonseca, A., Gavaret, M., Carron, R., Bartolomei, F., Liégeois-Chauvel, C., Régis, J., Chauvel, P., & Alario, F.-X. (2014). Simultaneous recording of MEG, EEG and intracerebral EEG during visual stimulation: from feasibility to single-trial analysis. NeuroImage, 99, 548–558.

Enatsu, R., & Mikuni, N. (2016). Invasive evaluations for epilepsy surgery: a review of the literature. Neurologia medico-chirurgica, 56(5), 221–227.

Fonov, V. S., Evans, A. C., McKinstry, R. C., Almli, C. R., & Collins, D. (2009). Unbiased nonlinear average age-appropriate brain templates from birth to adulthood. NeuroImage (47), S102.

Frauscher, B., Von Ellenrieder, N., Zelmann, R., Doležalová, I., Minotti, L., Olivier, A., Hall, J., Hoffmann, D., Nguyen, D. K., & Kahane, P. (2018). Atlas of the normal intracranial electroencephalogram: neurophysiological awake activity in different cortical areas. Brain, 141(4), 1130–1144.

Giraud, A.-L., & Poeppel, D. (2012). Cortical oscillations and speech processing: emerging computational principles and operations. Nature neuroscience, 15(4), 511–517.

Gramfort, A., Papadopoulo, T., Olivi, E., & Clerc, M. (2010). OpenMEEG: opensource software for quasistatic bioelectromagnetics. Biomedical engineering online, 9(1), 1–20.

Groppe, D. M., Bickel, S., Keller, C. J., Jain, S. K., Hwang, S. T., Harden, C., & Mehta, A. D. (2013). Dominant frequencies of resting human brain activity as measured by the electrocorticogram. NeuroImage, 79, 223–233.

Grova, C., Aiguabella, M., Zelmann, R., Lina, J. M., Hall, J. A., & Kobayashi, E. (2016). Intracranial EEG potentials estimated from MEG sources: A new approach to correlate MEG and iEEG data in epilepsy. Human brain mapping, 37(5), 1661–1683.

Grova, C., Daunizeau, J., Lina, J.-M., Bénar, C. G., Benali, H., & Gotman, J. (2006). Evaluation of EEG localization methods using realistic simulations of interictal spikes. NeuroImage, 29(3), 734–753.

Hamandi, K., Routley, B. C., Koelewijn, L., & Singh, K. D. (2016). Non-invasive brain mapping in epilepsy: applications from magnetoencephalography. Journal of Neuroscience Methods, 260, 283–291.

He, B. J. (2014). Scale-free brain activity: past, present, and future. Trends in cognitive sciences, 18(9), 480–487.

Hedrich, T., Pellegrino, G., Kobayashi, E., Lina, J.-M., & Grova, C. (2017). Comparison of the spatial resolution of source imaging techniques in high-density EEG and MEG. NeuroImage, 157, 531–544.

Heers, M., Chowdhury, R. A., Hedrich, T., Dubeau, F., Hall, J. A., Lina, J.-M., Grova, C., & Kobayashi, E. (2016). Localization accuracy of distributed inverse solutions for electric and magnetic source imaging of interictal epileptic discharges in patients with focal epilepsy. Brain topography, 29(1), 162–181.

Hillebrand, A., Barnes, G. R., Bosboom, J. L., Berendse, H. W., & Stam, C. J. (2012). Frequency-dependent functional connectivity within resting-state networks: an atlas-based MEG beamformer solution. NeuroImage, 59(4), 3909–3921.

Hipp, J. F., Hawellek, D. J., Corbetta, M., Siegel, M., & Engel, A. K. (2012). Large-scale cortical correlation structure of spontaneous oscillatory activity. Nature neuroscience, 15(6), 884–890.

Hirano, Y., & Uhlhaas, P. J. (2021). Current findings and perspectives on aberrant neural oscillations in schizophrenia. Psychiatry and clinical neurosciences, 75(12), 358–368.

Huang, Y., Sun, B., Debarros, J., Zhang, C., Zhan, S., Li, D., Zhang, C., Wang, T., Huang, P., & Lai, Y. (2021). Increased theta/alpha synchrony in the habenula-prefrontal network with negative emotional stimuli in human patients. Elife, 10.

Jayakar, P., Gotman, J., Harvey, A. S., Palmini, A., Tassi, L., Schomer, D., Dubeau, F., Bartolomei, F., Yu, A., & Kršek, P. (2016). Diagnostic utility of invasive EEG for epilepsy surgery: indications, modalities, and techniques. Epilepsia, 57(11), 1735–1747.

Kakisaka, Y., Kubota, Y., Wang, Z. I., Piao, Z., Mosher, J. C., Gonzalez-Martinez, J., Jin, K., Alexopoulos, A. V., & Burgess, R. C. (2012). Use of simultaneous depth and MEG recording may provide complementary information regarding the epileptogenic region. Epileptic disorders, 14(3), 298–303.

Keitel, A., & Gross, J. (2016). Individual human brain areas can be identified from their characteristic spectral activation fingerprints. PLoS biology, 14(6), e1002498.

Koessler, L., Benar, C., Maillard, L., Badier, J.-M., Vignal, J. P., Bartolomei, F., Chauvel, P., & Gavaret, M. (2010). Source localization of ictal epileptic activity investigated by high resolution EEG and validated by SEEG. NeuroImage, 51(2), 642–653.

Kybic, J., Clerc, M., Abboud, T., Faugeras, O., Keriven, R., & Papadopoulo, T. (2005). A common formalism for the integral formulations of the forward EEG problem. IEEE transactions on medical imaging, 24(1), 12–28.

Landman, B. A., & Warfield, S. K. (2019). MICCAI 2012: Workshop on multi-atlas labeling.éditeur non identifié.

Lapalme, E., Lina, J.-M., & Mattout, J. (2006). Data-driven parceling and entropic inference in MEG. NeuroImage, 30(1), 160–171.

Lina, J.-M., Chowdhury, R., Lemay, E., Kobayashi, E., & Grova, C. (2012). Wavelet-based localization of oscillatory sources from magnetoencephalography data. IEEE transactions on biomedical engineering, 61(8), 2350–2364.

Mahjoory, K., Schoffelen, J.-M., Keitel, A., & Gross, J. (2020). The frequency gradient of human restingstate brain oscillations follows cortical hierarchies. Elife, 9, e53715.

Mattout, J., Pélégrini-Issac, M., Garnero, L., & Benali, H. (2005). Multivariate source prelocalization (MSP): use of functionally informed basis functions for better conditioning the MEG inverse problem. NeuroImage, 26(2), 356–373.

Mellem, M. S., Wohltjen, S., Gotts, S. J., Ghuman, A. S., & Martin, A. (2017). Intrinsic frequency biases and profiles across human cortex. Journal of neurophysiology, 118(5), 2853–2864.

Niso, G., Rogers, C., Moreau, J. T., Chen, L.-Y., Madjar, C., Das, S., Bock, E., Tadel, F., Evans, A. C., & Jolicoeur, P. (2016). OMEGA: the open MEG archive. NeuroImage, 124, 1182–1187.

Niso, G., Tadel, F., Bock, E., Cousineau, M., Santos, A., & Baillet, S. (2019). Brainstorm pipeline analysis of resting-state data from the open MEG archive. Frontiers in neuroscience, 13, 284.

Nunez, P. L., Wingeier, B. M., & Silberstein, R. B. (2001). Spatial-temporal structures of human alpha rhythms: Theory, microcurrent sources, multiscale measurements, and global binding of local networks. Human brain mapping, 13(3), 125–164.

Ostlund, B. D., Alperin, B. R., Drew, T., & Karalunas, S. L. (2021). Behavioral and cognitive correlates of the aperiodic (1/f-like) exponent of the EEG power spectrum in adolescents with and without ADHD. Developmental cognitive neuroscience, 48, 100931.

Ouyang, G., Hildebrandt, A., Schmitz, F., & Herrmann, C. S. (2020). Decomposing alpha and 1/f brain activities reveals their differential associations with cognitive processing speed. NeuroImage, 205, 116304.

Pellegrino, G., Hedrich, T., Chowdhury, R., Hall, J. A., Lina, J. M., Dubeau, F., Kobayashi, E., & Grova, C. (2016). Source localization of the seizure onset zone from ictal EEG/MEG data. Human brain mapping, 37(7), 2528–2546.

Pellegrino, G., Hedrich, T., Chowdhury, R. A., Hall, J. A., Dubeau, F., Lina, J. M., Kobayashi, E., & Grova, C. (2018). Clinical yield of magnetoencephalography distributed source imaging in epilepsy: A comparison with equivalent current dipole method. Human brain mapping, 39(1), 218–231.

Pellegrino, G., Hedrich, T., Sziklas, V., Lina, J. M., Grova, C., & Kobayashi, E. (2021). How cerebral cortex protects itself from interictal spikes: The alpha/beta inhibition mechanism. Human brain mapping, 42(11), 3352–3365.

Pellegrino, G., Schuler, A.-L., Arcara, G., Di Pino, G., Piccione, F., & Kobayashi, E. (2022). Resting state network connectivity is attenuated by fMRI acoustic noise. NeuroImage, 247, 118791.

Pellegrino, G., Schuler, A. L., Arcara, G., Di Pino, G., Piccione, F., & Kobayashi, E. (2022). Resting state network connectivity is attenuated by fMRI acoustic noise. NeuroImage 247, 118791.

Penfield, W., & Jasper, H. (1954). Epilepsy and the functional anatomy of the human brain.

Pizzo, F., Roehri, N., Medina Villalon, S., Trébuchon, A., Chen, S., Lagarde, S., Carron, R., Gavaret, M., Giusiano, B., & McGonigal, A. (2019). Deep brain activities can be detected with magnetoencephalography. Nature communications, 10(1), 1–13.

Prichard, D., & Theiler, J. (1994). Generating surrogate data for time series with several simultaneously measured variables. Physical review letters, 73(7), 951.

Rampp, S., Kaltenhäuser, M., Weigel, D., Buchfelder, M., Blümcke, I., Dörfler, A., & Stefan, H. (2010). MEG correlates of epileptic high gamma oscillations in invasive EEG. Epilepsia, 51(8), 1638–1642.

Ramsay, I. S., Lynn, P. A., Schermitzler, B., & Sponheim, S. R. (2021). Individual alpha peak frequency is slower in schizophrenia and related to deficits in visual perception and cognition. Scientific reports, 11(1), 1–9.

Santiuste, M., Nowak, R., Russi, A., Tarancon, T., Oliver, B., Ayats, E., Scheler, G., & Graetz, G. (2008). Simultaneous magnetoencephalography and intracranial EEG registration: technical and clinical aspects. Journal of Clinical Neurophysiology, 25(6), 331–339.

Schaworonkow, N., & Voytek, B. (2021). Longitudinal changes in aperiodic and periodic activity in electrophysiological recordings in the first seven months of life. Developmental cognitive neuroscience, 47, 100895.

Schnitzler, A., & Gross, J. (2005). Normal and pathological oscillatory communication in the brain. Nature reviews neuroscience, 6(4), 285–296.

Senoussi, M., Verbeke, P., Desender, K., De Loof, E., Talsma, D., & Verguts, T. (2022). Theta oscillations shift towards optimal frequency for cognitive control. Nature Human Behaviour, 1–14.

Srinivasan, R., Winter, W. R., & Nunez, P. L. (2006). Source analysis of EEG oscillations using high-resolution EEG and MEG. Progress in brain research, 159, 29–42.

Tadel, F., Baillet, S., Mosher, J. C., Pantazis, D., & Leahy, R. M. (2011). Brainstorm: a user-friendly application for MEG/EEG analysis. Computational intelligence and neuroscience, 2011.

Uusitalo, M. A., & Ilmoniemi, R. J. (1997). Signal-space projection method for separating MEG or EEG into components. Medical and biological engineering and computing, 35(2), 135–140.

von Ellenrieder, N., Beltrachini, L., & Muravchik, C. H. (2012). Electrode and brain modeling in stereo-EEG. Clinical Neurophysiology, 123(9), 1745–1754.

von Ellenrieder, N., Khoo, H. M., Dubeau, F., & Gotman, J. (2021). What do intracerebral electrodes measure? Clinical Neurophysiology, 132(5), 1105–1115.

von Ellenrieder, N., Pellegrino, G., Hedrich, T., Gotman, J., Lina, J.-M., Grova, C., & Kobayashi, E. (2016). Detection and magnetic source imaging of fast oscillations (40–160 Hz) recorded with magnetoencephalography in focal epilepsy patients. Brain topography, 29(2), 218–231.

Voytek, B., Canolty, R. T., Shestyuk, A., Crone, N. E., Parvizi, J., & Knight, R. T. (2010). Shifts in gamma phase–amplitude coupling frequency from theta to alpha over posterior cortex during visual tasks. Frontiers in human neuroscience, 4, 191.

Wang, X.-J. (2010). Neurophysiological and computational principles of cortical rhythms in cognition. Physiological reviews, 90(3), 1195–1268.

Wen, H., & Liu, Z. (2016). Separating fractal and oscillatory components in the power spectrum of neurophysiological signal. Brain topography, 29(1), 13–26.

Wiesman, A. I., da Silva Castanheira, J., & Baillet, S. (2022). Stability of spectral estimates in resting-state magnetoencephalography: Recommendations for minimal data duration with neuroanatomical specificity. NeuroImage, 247, 118823.

Wilkinson, C. L., & Nelson, C. A. (2021). Increased aperiodic gamma power in young boys with Fragile X Syndrome is associated with better language ability. Molecular autism, 12(1), 1–15.

Zhang, Y., Van Drongelen, W., & He, B. (2006). Estimation of in vivo brain-to-skull conductivity ratio in humans. Applied physics letters, 89(22), 223903.

