## supplementary material for "Validating MEG source imaging of resting state oscillatory patterns with an intracranial EEG atlas"

### MEG relative power before and after conversion to IEEG space

Fig S2 confirms that after conversion from MEG source map to intracranial space (VIEEG), the activity in virtual intracranial space (VIEEG) was concordant with the MEG source activity along the cortical surface. The MEG relative power along the cortical surface for different frequency bands was actually reproducing the spatial distribution of relative power maps reported by previous MEG studies using MNE localization (Niso et al., 2016; Niso et al., 2019).

MEG estimated spectra showed strongest delta activity along the gyrus rectus and orbital gyrus region. MEG estimated delta power in hippocampus was also present (Fig S1). Our MEG estimated delta activity was consistent with the previous findings, reporting strong delta activity along hippocampal areas and orbitofrontal regions (Congedo et al., 2010; Congedo, 2010; Hillebrand et al., 2012; Niso et al., 2016; Niso et al., 2019).

Theta activity was previously reported to be centered over midline fronto-central regions (Scheeringa et al., 2008), dorsal anterior cingulate cortex (ACC) or medial prefrontal cortex (Gevins et al., 1997; Onton et al., 2005), dorsal frontal cortex (Kalamangalam et al., 2020; Mellem et al., 2017; Niso et al., 2019). Our MEG results in theta band were also localized in these regions and consistent with previous MEG studies.

Relative power in alpha band was strongest in occipital and parietal areas. Compared to other brain regions in alpha band, low alpha power was found in frontal regions. Our findings are consistent with previous studies (Niso et al., 2019, Niso et al., 2016), suggesting strongest alpha activity in precuneus (Capilla et al., 2022; Hari & Salmelin, 1997; Salmelin & Hari, 1994) and strong activity over visual, auditory, and somatosensory regions (Haegens et al., 2015; Hari & Salmelin, 1997; Salmelin & Hari, 1994). The widespread relative power in alpha band estimated by MEG was consistent with the activity found in (Capilla et al., 2022), where they reported widespread dominance in alpha peak along the posterior regions.

Previous studies reported beta activity in motor area (Frauscher et al., 2018; Hillebrand et al., 2012; Jasper & Penfield, 1949; Niso et al., 2016) and prefrontal areas (Mahjoory et al., 2020; Mellem et al., 2017). Consistent with such findings, we also found strong beta relative power in motor areas and prefrontal regions. However, we found that the strongest beta relative power estimated by wMEM was in the lateral parietal regions (Fig S1). In the gamma band, the frontal regions showed overall strong activity. Frontal gamma activity in MEG was also reported in previous studies (Niso et al., 2016; Niso et al., 2019).

In Fig S2, we compared wMEM and depth weighted MNE in the intracranial IEEG space and along the cortical surface. It is important to note that the color bar is ranging from maximum to minimum value in each modality, and due to the inherent nature of wMEM method, some sources are shut down resulting in zero activity within those regions. Therefore, wMEM resulting maps along the cortical surface are ranging from zero to maximum. Overall, both source imaging methods are generating similar maps of relative power, with differences mostly in the gamma band, where MNE localization was more anterior and of larger amplitude when compared to wMEM. In the beta band, wMEM localization was more posterior compared to MNE.

### Supplementary figures:

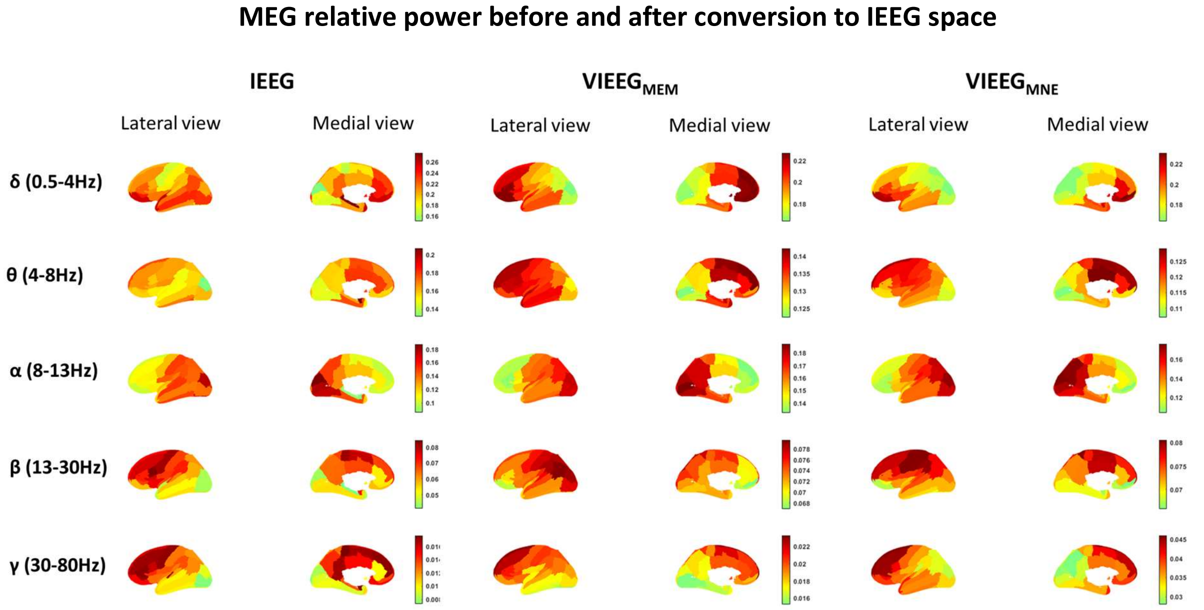


Figure S1: Group average of relative PSD values across each frequency band and over all the available channels in each ROI of the IEEG atlas, the ground truth, the MEG estimated VIEEG using wavelet-MEM (wMEM) method $VIEEG_{MEM}$, and minimum norm method estimate $VIEEG_{MNE}$. The number of channels in each ROI ($N_{ROI}$) in IEEG varies. For each ROI, VIEEG was estimated for $45 \times N_{ROI}$ channels, where total number of subjects is 45. The relative PSD for each channel is calculated as the ratio of the power of the signal in each frequency bin relatively to the total power of the signal. Relative PSD value for a channel range between 0 and 1. The color bar ranges from minimum to maximum value for each modality in each frequency band.

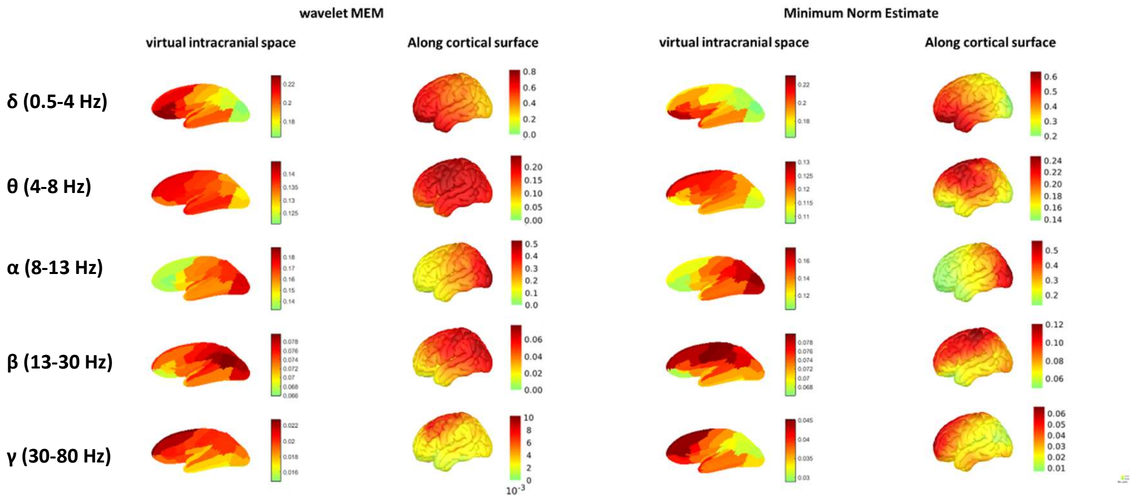


Figure S2: Comparison of group average of relative PSD values between MEG source imaging along the cortical surface and MEG estimated VIEEG, for two source imaging methods: (A) wavelet-MEM (wMEM) and (B) minimum norm method estimate. The relative PSD for each channel in VIEEG or each vertex along the cortical surface is calculated as the ratio of the power of the signal in each frequency bin relatively to the total power of the signal. Relative PSD value ranges between 0 and 1. The color bar ranges from minimum to maximum value for each modality and in each frequency band.

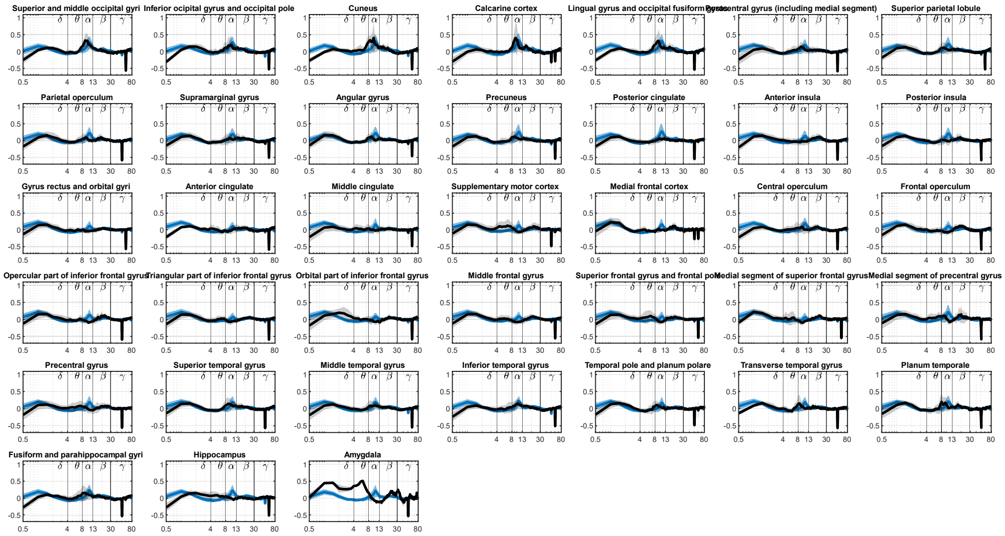


Figure S3: Normalized PSD after removing aperiodic components for all 38 ROIs for IEEG and VIEEG.

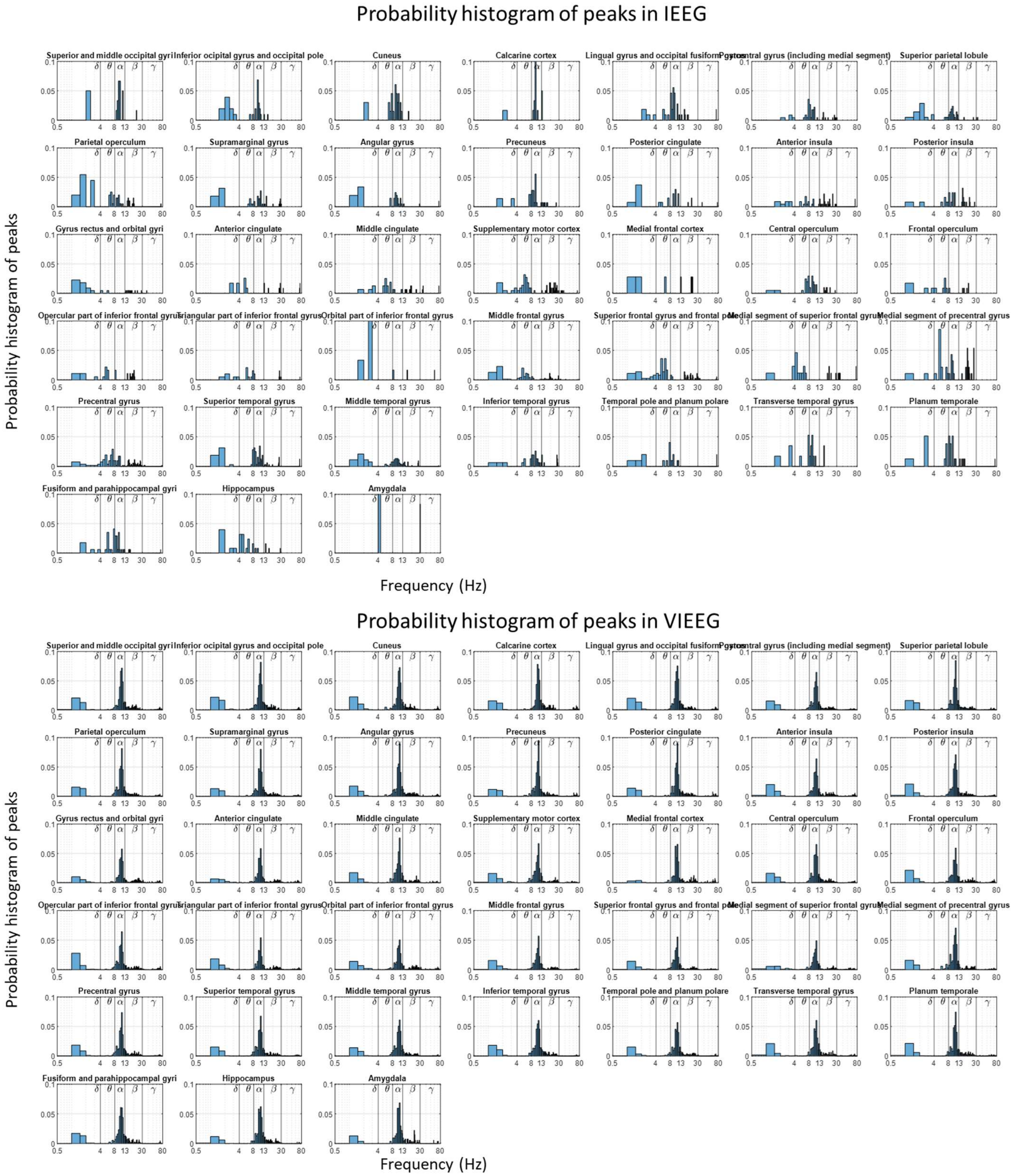


Figure S4: Probability histogram of all identified peaks in each ROI plotted for all 38 ROIs for IEEG and $V_{IEEG}$

### Comparing common-average versus bipolar montage

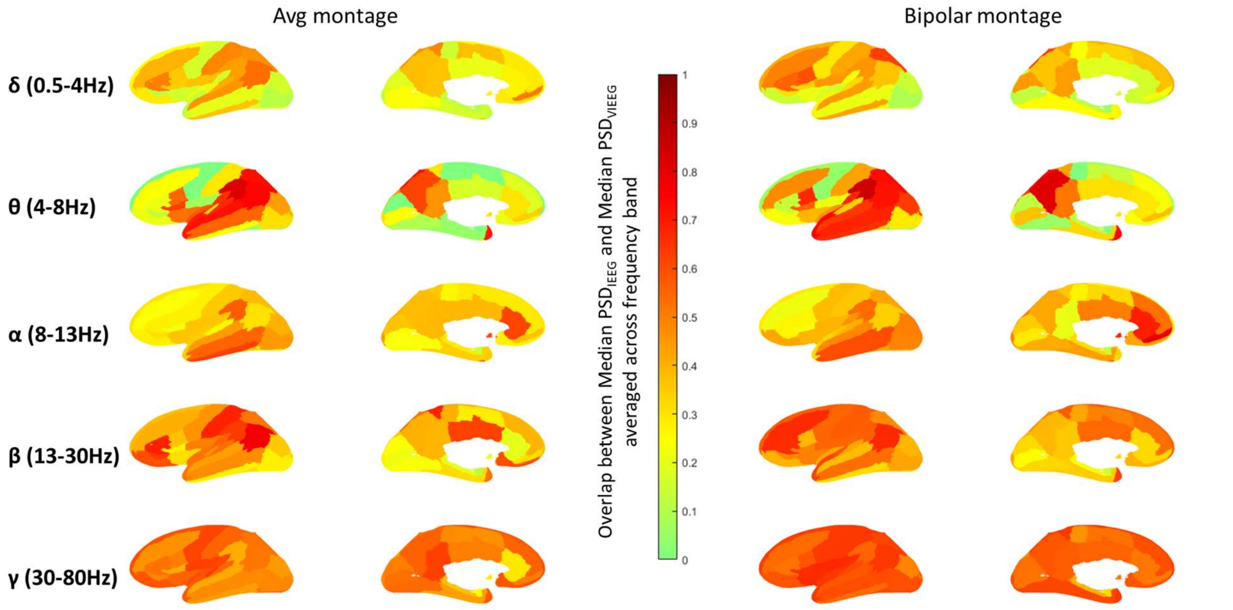


Figure S5: *Average overlap* between VIEEG and IEEG spectra across each spectral band for each of the 38 ROIs for common average montage and bipolar montage. The value of *overlap* is calculated at each frequency bin and ranges between 0 to 1. For a ROI, if the median of $PSD_{VIEEG}$ perfectly coincides with median of $PSD_{IEEG}$ at all frequency bins within a specific frequency band, the average *overlap* is 1

### Comparing peak estimated by MEG: common-average versus bipolar montage

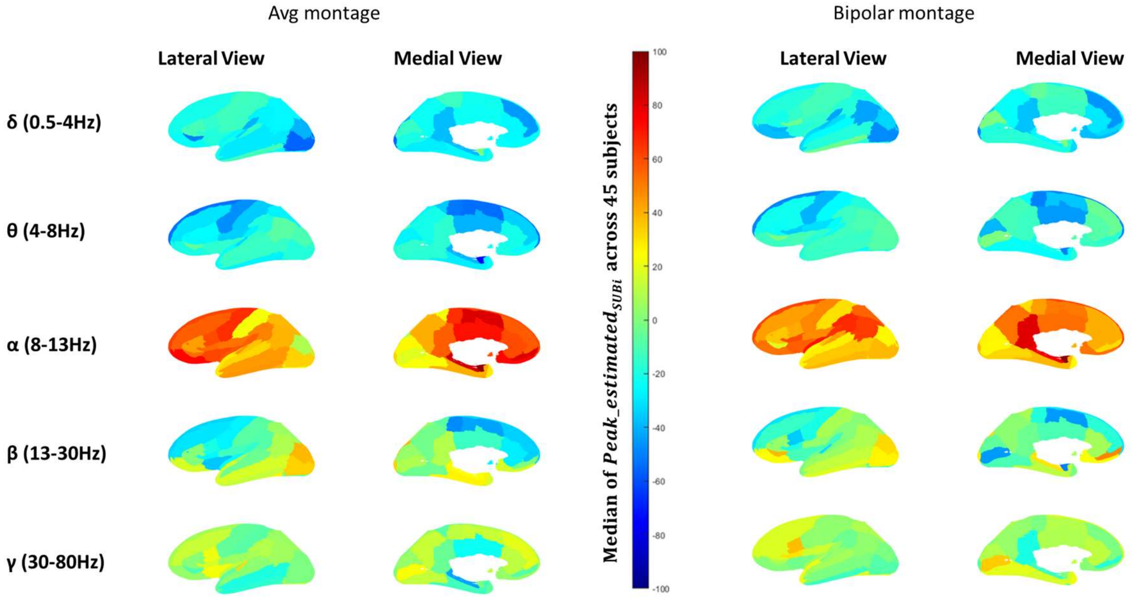


Figure S6: Comparison of median value of $Peak\_estimated_{SUBi}$ over 45 subjects between common average montage and bipolar montage. $Peak\_estimated_{SUBi}$ measures the percentage difference of number of channels exhibiting spectral peaks in VIEEG compared to IEEG, as a proportion of the total number of channels in each ROI, for each spectral band and ROI, calculated for each subject *i*.

### References:


Capilla, A., Arana, L., García-Huéscar, M., Melcón, M., Gross, J., & Campo, P. (2022). The natural frequencies of the resting human brain: an MEG-based atlas. *NeuroImage*, 119373.

Congedo, M., John, R. E., De Ridder, D., & Prichep, L. (2010). Group independent component analysis of resting state EEG in large normative samples. *International Journal of Psychophysiology*, *78*(2), 89-99.

Congedo, M., Roy E. John, Dirk De Ridder, and Leslie Prichep. (2010). Group independent component analysis of resting state EEG in large normative samples. *International Journal of Psychophysiology 78, no. 2*, 89-99.

Frauscher, B., Von Ellenrieder, N., Zelmann, R., Doležalová, I., Minotti, L., Olivier, A., Hall, J., Hoffmann, D., Nguyen, D. K., & Kahane, P. (2018). Atlas of the normal intracranial electroencephalogram: neurophysiological awake activity in different cortical areas. *Brain*, *141*(4), 1130-1144.

Gevins, A., Smith, M. E., McEvoy, L., & Yu, D. (1997). High-resolution EEG mapping of cortical activation related to working memory: effects of task difficulty, type of processing, and practice. *Cerebral cortex (New York, NY: 1991)*, *7*(4), 374-385.

Haegens, S., Barczak, A., Musacchia, G., Lipton, M. L., Mehta, A. D., Lakatos, P., & Schroeder, C. E. (2015). Laminar profile and physiology of the α rhythm in primary visual, auditory, and somatosensory regions of neocortex. *Journal of Neuroscience*, *35*(42), 14341-14352.

Hari, R., & Salmelin, R. (1997). Human cortical oscillations: a neuromagnetic view through the skull. *Trends in neurosciences*, *20*(1), 44-49.

Hillebrand, A., Barnes, G. R., Bosboom, J. L., Berendse, H. W., & Stam, C. J. (2012). Frequency-dependent functional connectivity within resting-state networks: an atlas-based MEG beamformer solution. *NeuroImage*, *59*(4), 3909-3921.

Jasper, H., & Penfield, W. (1949). Electrocorticograms in man: effect of voluntary movement upon the electrical activity of the precentral gyrus. *Archiv für Psychiatrie und Nervenkrankheiten*, *183*(1), 163-174.

Kalamangalam, G. P., Long, S., & Chelaru, M. I. (2020). A neurophysiological brain map: Spectral parameterization of the human intracranial electroencephalogram. *Clinical Neurophysiology*, *131*(3), 665-675.

Mahjoory, K., Schoffelen, J.-M., Keitel, A., & Gross, J. (2020). The frequency gradient of human resting-state brain oscillations follows cortical hierarchies. *Elife*, *9*, e53715.

Mellem, M. S., Wohltjen, S., Gotts, S. J., Ghuman, A. S., & Martin, A. (2017). Intrinsic frequency biases and profiles across human cortex. *Journal of neurophysiology*, *118*(5), 2853-2864.

Niso, G., Rogers, C., Moreau, J. T., Chen, L.-Y., Madjar, C., Das, S., Bock, E., Tadel, F., Evans, A. C., & Jolicoeur, P. (2016). OMEGA: the open MEG archive. *NeuroImage*, *124*, 1182-1187.

Niso, G., Tadel, F., Bock, E., Cousineau, M., Santos, A., & Baillet, S. (2019). Brainstorm pipeline analysis of resting-state data from the open MEG archive. *Frontiers in neuroscience*, *13*, 284.

Onton, J., Delorme, A., & Makeig, S. (2005). Frontal midline EEG dynamics during working memory. *NeuroImage*, *27*(2), 341-356.

Salmelin, R., & Hari, R. (1994). Characterization of spontaneous MEG rhythms in healthy adults. *Electroencephalography and clinical neurophysiology*, *91*(4), 237-248.

Scheeringa, R., Bastiaansen, M. C., Petersson, K. M., Oostenveld, R., Norris, D. G., & Hagoort, P. (2008). Frontal theta EEG activity correlates negatively with the default mode network in resting state. *International Journal of Psychophysiology*, *67*(3), 242-251.
